# A simulation-based deep learning framework for spatially explicit malaria modeling of CRISPR suppression gene drive mosquitoes

**DOI:** 10.1101/2025.03.10.642387

**Authors:** Yuan Hu Allegretti, Weitang Sun, Jackson Champer

**Affiliations:** Center for Bioinformatics, School of Life Sciences, Center for Life Sciences, Peking University; Institute for AI Industry Research, Tsinghua University

**Author notes:** Equal contribution, authors may rearrange the order of names for references (Yuan Hu Allegretti, Weitang Sun).

## Abstract

Engineered CRISPR gene drives are a promising new strategy for fighting malaria and other vector-borne diseases, made possible by genome engineering with the CRISPR-Cas9 system. One useful approach to predict the outcome of a gene drive mosquito release is individual-based modeling, which can be spatially explicit and allows flexible parameters for drive efficiency, mosquito ecology, and malaria transmission. However, the computational demand of this type of model significantly increases when including a larger number of parameters, especially due to the chasing phenomenon, which can delay or prevent successful population elimination. Thus, we built a simulation-based deep-learning model to comprehensively understand the effects of different parameters on *Anopheles gambiae* mosquito suppression and malaria prevalence among the human population. The results suggest that reducing the embryo resistance cut rate, reducing the functional resistance forming rate and increasing the drive conversion rate plays the major role in mosquito suppression and related phenomena. We also observed that the parameter space for eliminating malaria was substantially larger than that for mosquito suppression, suggesting that even a considerably imperfect drive may still successfully accomplish its objective despite chasing or resistance allele formation. This study shows that suppression gene drives may be highly effective at locally eliminating malaria, even in challenging conditions.

## Introduction

Eradication of malaria is one of the most important public health objectives today. The World Health Organization (WHO) reports that there were 263 million cases and 597,000 deaths in 83 malaria endemic countries in 2023(Organization, 2024). The ongoing campaign to eradicate malaria includes distribution of personal protection, research and development of rapid malaria diagnosis methods, and novel malaria drugs and other measures(Group, 2006). However, the progress of the malaria eradication campaign has stalled since 2010 because the parasite and mosquitoes have developed resistance to frontline drugs and insecticides(Blasco et al., 2017; Ranson et al., 2009). Thus, development and deployment of new tools is needed for prevention and control of malaria.

Engineered CRISPR gene drives are a promising new strategy for fighting against malaria and other vector-borne diseases, made possible by the establishment of genome engineering with the CRISPR-Cas9 system(Hay et al., 2021; Verkuijl et al., 2022; Wang et al., 2022; Wang et al., 2024). Homing gene drives are genetic constructs that contain an endonuclease, such as Cas9 directed by a guide RNA (gRNA), to target a specific site in the genome for cleavage. After cleaving this target site, the construct then copies itself to that site via homology-directed repair, a process referred to as “homing” or “drive conversion”. This results in copying of the drive allele into the wild-type chromosome, converting the drive heterozygote into a homozygote in the germline. Because of super-Mendelian inheritance, a gene drives can rapidly spread in the target population(Burt, 2003; Deredec et al., 2008). Such gene drives can be used for population modification or for suppression.

Suppression gene drives usually target a haplosufficient but essential female fertility gene, eventually leading to population reduction due to sterile female homozygotes. In addition to the suppression of the target species, some targets such as the female-specific exon of *doublesex* result in intersex sterile female homozygous mosquitoes that do not take blood meals, reducing disease spread more quickly(Kyrou et al., 2018). Early suppression drives in *Anopheles* mosquitoes were sensitive to functional resistance allele formation and female fitness costs(Hammond et al., 2016), but more recent designs may be able to overcome these obstacles using techniques such as multiplexed gRNAs(Du et al., 2024; Xu et al., 2025), conserved target sites(Kyrou et al., 2018), and improved Cas9 promoters(Du et al., 2024; Hammond et al., 2021), thus increasing prospects for field deployment.

While there is growing evidence showing that suppression gene drives are promising tools to control mosquito populations(Champer et al., 2022; Eckhoff et al., 2017; Mondal et al., 2024a; North et al., 2020; Sánchez C et al., 2020; Wu et al., 2021), little is known about the effect of suppression gene drive on malaria prevalence(Eckhoff et al., 2017). This is particularly important because recent studies indicate that the “chasing” phenomenon (wherein wild-type escapes to empty space and avoid the drive, but the drive continues to “chase” wild-type in an arena) may prevent successful population elimination by a drive, even if the drive can eliminate panmictic populations(Birand et al., 2022; J. Champer et al., 2021; Liu & Champer, 2022; Liu et al., 2023; Paril & Phillips, 2022; Zhu & Champer, 2023). Such chasing can also eventually lead to functional resistance allele formation(J. Champer et al., 2021; Liu & Champer, 2022). A recent study used a patch-based model to estimate the effect of suppression drive on malaria transmission with promising results(Hancock et al., 2024) and other patch based models for gene drive can also simulate disease(Mondal et al., 2024a). However, it is unclear if mosquito population structures are better represented by networks of connected patches(Mondal et al., 2024a; Sánchez C et al., 2020; Wu et al., 2021) or continuous space (as in this study). Both may be most suitable for different scales. This study also implemented whole-arena malaria rates based on biting female population size, preventing disease rates from varying over space, which may produce different results.

Here, we aimed to model how a suppression gene drive could affect rates of malaria in a human population threatened by *Anopheles gambiae*. However, due to the large number of parameters for disease, mosquito ecology(Dhole et al., 2020; Kim et al., 2023), and gene-drive performance, performing simulations while systematically varying all parameters would have a high computational burden, also known as the “curse of dimensionality”(Berisha et al., 2021). Therefore, we built a deep-learning based model to comprehensively understand the role of each parameter of interest. The deep learning algorithm automatically selects features that have a high impact on the outcome, enabling us to search the full parameter space with a low computational cost without fixing any parameters(Poggio et al., 2017). With our model, we then predicted the possibility of successful mosquito and malaria elimination. We found that current gene drives are likely sufficient to significantly reduce the prevalence of malaria in a target population and that the parameter space of malaria eradication is significantly larger than that for mosquito eradication. The chasing phenomenon may positively contribute to malaria elimination despite preventing mosquito population elimination, showcasing the important of spatiality in predicting disease transmission. This study shows the potential to utilize simulation-based deep learning models to predict the impact of epidemiological interventions on vector-borne disease.

## Methods

### 1. Population model

#### 1.1 Gene drive mechanisms

We model a homing population suppression gene drive targeting an essential but haplosufficient female fertility gene. With a germline specific promoter for Cas9, the homing drive induces double strain breaks at its target site in germline cells. Through homology-directed repair, wild-type alleles will be converted to drive alleles at a rate equal to the drive conversion efficiency. Because the target gene is haplosufficient, drive/wild type heterozygote females remain fertile. When the drive allele reaches a high frequency in the population, there will be many sterile female drive homozygotes because these females lack any functional copies of the essential target gene. With few fertile females, the population may decline or even be eliminated.

However, double-stranded breaks in DNA can also be fixed by end joining repair. This pathway will often cause indel mutations in the target site, which may prevent recognition by the drive’s gRNAs. For gene drive, these mutated sequences are called resistance alleles and form at the germline resistance allele formation rate. They are resistant to drive cleavage and can’t be converted to drive alleles. Most of these are nonfunctional resistance alleles. These also cause recessive female sterility, but they can slow down the speed of the drive and reduce its equilibrium frequency(Beaghton et al., 2019). Functional resistance alleles can also form, albeit at much lower frequency (the relative functional resistance allele formation rate). However, they have a major fitness advantage over the drive and will tend to rapidly increase in frequency and eliminate the drive. Aside from the germline, resistance alleles can also form in early embryos from drive-carrying mothers at the embryo resistance allele formation rate. These immediately change the genotype of the offspring.

Although female drive heterozygotes are potentially fully fertile due to the haplosufficiency of the target gene, somatic expression of Cas9 can reduce the fertility of female drive heterozygotes(Hammond et al., 2016; Kyrou et al., 2018; Xu et al., 2024; Yang et al., 2022). It is also possible for drive fitness costs to reduce drive offspring viability. This type of fitness cost is multiplicative per drive allele and applies to both sexes.

1.2 Mosquito model

We used a SLiM 4 model(Haller & Messer, 2023) as described in our previous study as the basis for our new model(Champer et al., 2022). We simulate the life history of *Anopheles* mosquitoes using one week time steps, during which we assume that the mosquitoes can complete one reproductive cycle. At the start of every cycle, adult females perform the following steps in sequence: finding a mate, searching for a blood meal, and generating offspring. Female mosquitos often mate only once, so if they have already found a mate, there is only a 5% chance that they will remate each week, and we assume that all eggs are fertilized by the most recent one (Frédéric, 2003).

When finding a mate, females choose a random male within the average dispersal distance (Figure S1). They will use this genotype as the father to generate offspring in the current week and in future weeks. If a female has already mated (either in the current week or a previous week), it has a certain probability to search for a blood meal. It will bite a random human within the dispersal distance (Figure S1) (*A. gambiae* females specialize in finding and biting humans), but if this is not possible, it will have a fixed probability to find and bite an animal, thus allowing it to reproduce without any possibility of malaria transmission. The number of offspring follows poison distribution with an average of 36. Real female mosquitoes tend to lay many more eggs after a single blood meal, but such offspring will usually have high mortality even in ideal natural conditions, so we choose a number that was high enough to allow some variability in surviving offspring number even when larval competition was very low (resulting in maximum larval survival). Drive/wild-type heterozygote females may have fewer offspring if they have fitness costs due to somatic expression of Cas9 and gRNA.

The survival rate of new age 0 mosquitoes (larvae) is controlled by the local density and their own genotype-based fitness. Old larvae have a relative high competition coefficient against new larvae. This coefficient was set to 5 as a default. To gain better runtime performance, a resource-explicit interaction is used during the larva competition calculation(Champer et al., 2024). Hexagonal tiling is utilized with an inelastic model(Champer et al., 2024). The foraging radius of larvae is set to 33 m. One foraging area contains 12 resources nodes on average. In each resource node, newly hatched larvae suffer competition pressure.

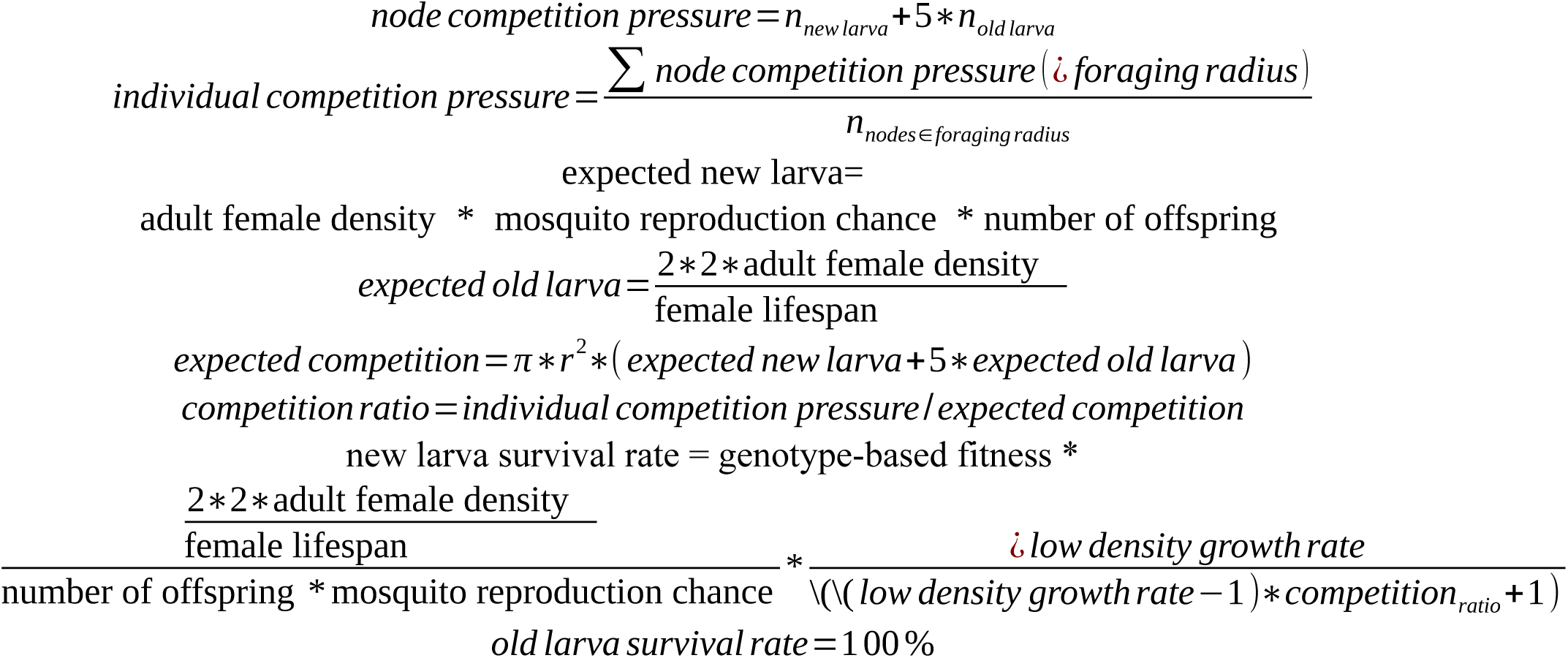

For adults, we use linear survival curves. Females live at most 7 weeks and males at most 5 weeks according to the equations:

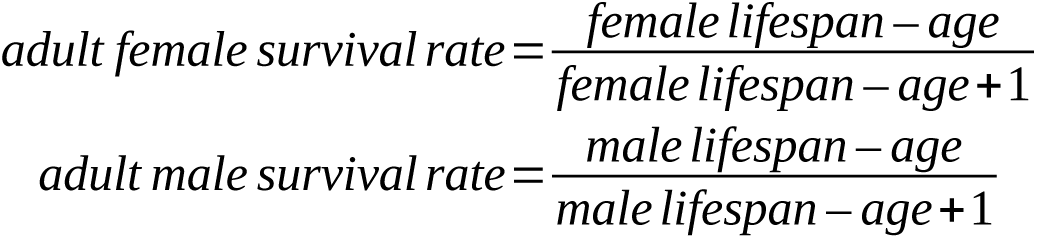

1.3 Spatial model

Every simulation takes place at a 4 km x 4 km arena. We set human density to 300 people per square kilometer, representing intensively cultivated farmland. Our mosquito density of 3000 adult females per square kilometer represents the effective population size of these(Messina et al., 2011). Initial human and mosquito placement is random. Humans do not move during the simulation, representing the fact that many *A. gambiae* bites take place at night, and most people rest in the same house every day. Human reproduction and death are not modeled. *A. gambiae* often reproduce in small, temporary habitats(Minakawa et al., 2004), so larvae are not able to travel far. We thus do not model any mosquito movement until they become adults after two weeks. Adults displace at the end of every week. Assuming *Anopheles* are moving randomly in every time step, we model a normally distribution in a random direction with a parameter representing the average dispersal. New larvae are displaced in an identical manner from their mother. Movement beyond the simulated area is not allowed, so positions are redrawn if any individuals are moved out of the arena.

1.4 Malaria transmission model

We model transmission of the malaria parasite *Plasmodium falciparum* between humans and mosquitoes. To simplify the model, we assume that *Anopheles gambiae* is the only vector in the area that carries malaria, and that there are no animal reservoirs of malaria. Humans and mosquitoes are represented as individuals, and malaria is represented as a status in individuals. Details of the model are shown in Figure 1. The constants in the model are shown in Table S1.

**Figure 1.**
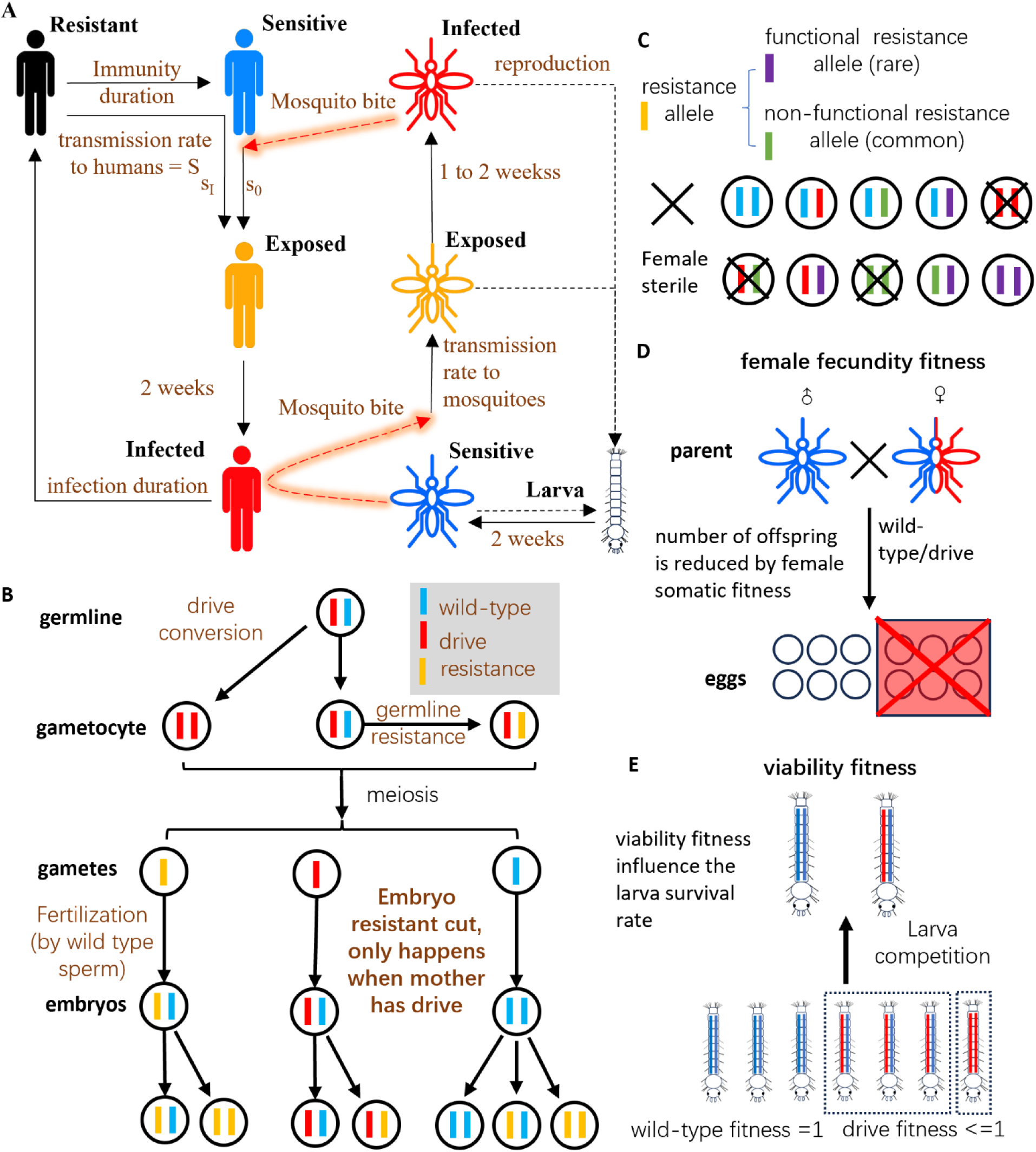
Model diagram. **A** Malaria transmission flow diagram for the human-malaria-mosquito model. Humans are classified into four infection states. Parameters include: s_0_, transmission rate to naïve humans. s_I_, transmission rate to resistant humans. Malaria transmission between humans and mosquitoes is highlighted in orange. All newly hatched mosquitoes do not carry malaria. Colors represent different infection states. **B** Gene drive inheritance in the mosquitoes. The homing gene drive functions in the germline drive/wild-type heterozygotes. Cas9 forms double-stranded breaks at the wild-type allele. If homology-directed repair occurs, the wild-type allele will be converted into a drive allele. Instead of drive conversion, resistance alleles could form by end-joining. Resistance alleles can also form in the early embryo through maternal deposition of Cas9 and gRNA. **C** Resistance alleles include functional and non-functional versions. Only females with at least one copy of a wild-type allele or functional resistance allele are fertile. **D** The progeny number of drive heterozygote females can be reduced due to fitness costs from somatic Cas9 expression. **E** Other fitness costs can reduce the viability of drive carriers.

Humans are divided into four stages according to their malaria infection status. They are Sensitive, Exposed, Infectious, or Resistant. Sensitive humans have a fixed chance ( *s*_0_ ) to become Exposed after being bitten by an infectious *Anopheles* mosquito. Exposed humans becomes Infectious two weeks later based on the *Plasmodium falciparum* exo-erythrocytic stage of about 6 days, which then requires 7-15 days to generate gametes from the erythrocytic asexual stage(Talman et al., 2004). Humans in the Infectious stage can transmit disease to *Anopheles* if they are bitten. It takes about 6 to 21 weeks to recover from malaria(Falk et al., 2006; Griffin et al., 2010; Macdonald. G., 1957; McKenzie & Bossert, 2005; Sama et al., 2005), so we assume that duration follows a Poisson distribution with a variable average. Humans in the Resistant stage have a lower infection rate depending on their immunity. The sensitivity of malaria infection for each human in the resistant stage is:

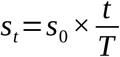

In this function, *t* is the time after recovery, and *T* represents the maximum duration of immunity, which declines linearly over time. Humans then become sensitive again after time *T*. *T* is determined individually for each infection and follows Poisson distribution with a specified average parameter.

Female Anopheles search for potential hosts at a frequency of *f _b_* every week. When a naive *Anopheles* bites an infectious human(Gu et al., 2003), it has a probability of 0.05 to 0.27(0.11 as default) becoming exposed. *Plasmodium falciparum* gametes need time to mature in mosquito, it takes one week for 40% of mosquitoes and two weeks for the rest(Shaw et al., 2020). After this period, the mosquitoes can transmit malaria to humans. *Anopheles* that are infected will retain their infection status indefinitely.

1. 2. **Deep learning framework**

2.1 Parameters and sampling

We trained a deep learning model on our SLiM simulation model. The variable parameters and their ranges are listed in Table 1. These were selected based on our previous findings(Champer et al., 2022) and literature search(Falk et al., 2006; Griffin et al., 2010; Gu et al., 2003; Macdonald. G., 1957; McKenzie & Bossert, 2005; Sama et al., 2005; Shaw et al., 2020; Talman et al., 2004). For parameters with little data available, we set a wide to ensure that realistic values would be contained within the range.

**Table 1.**
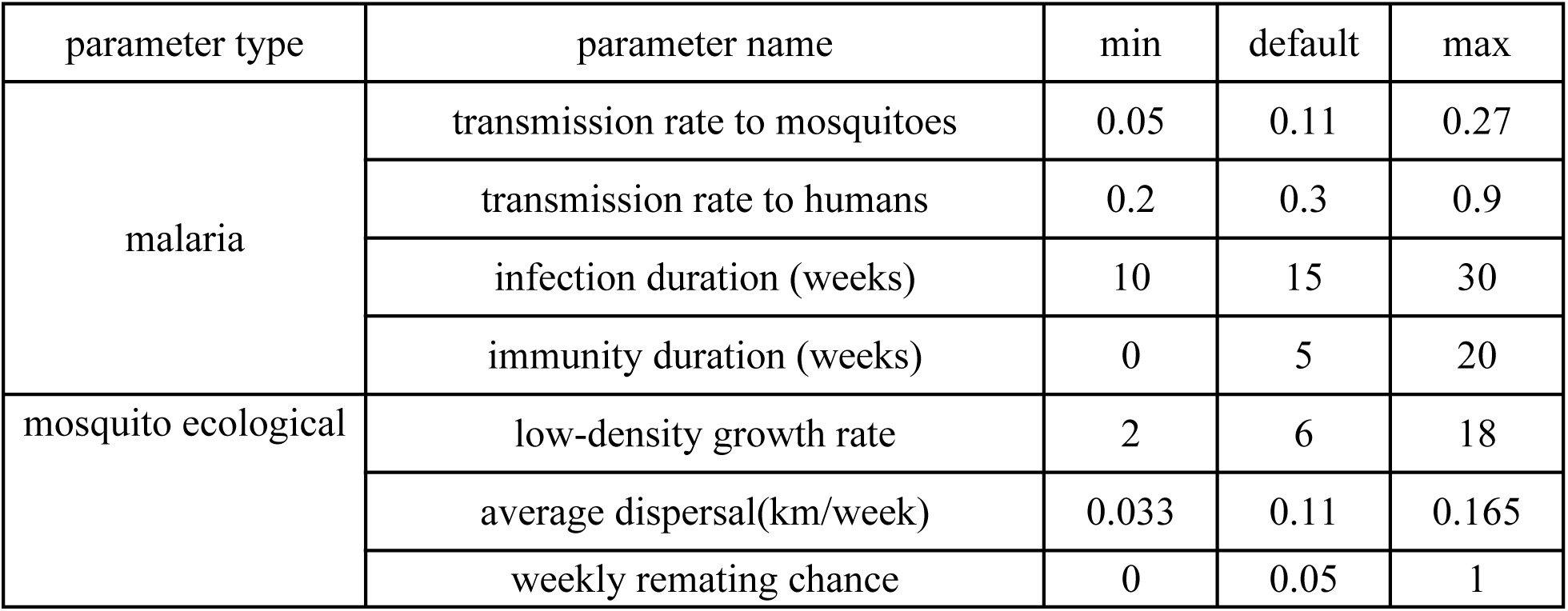
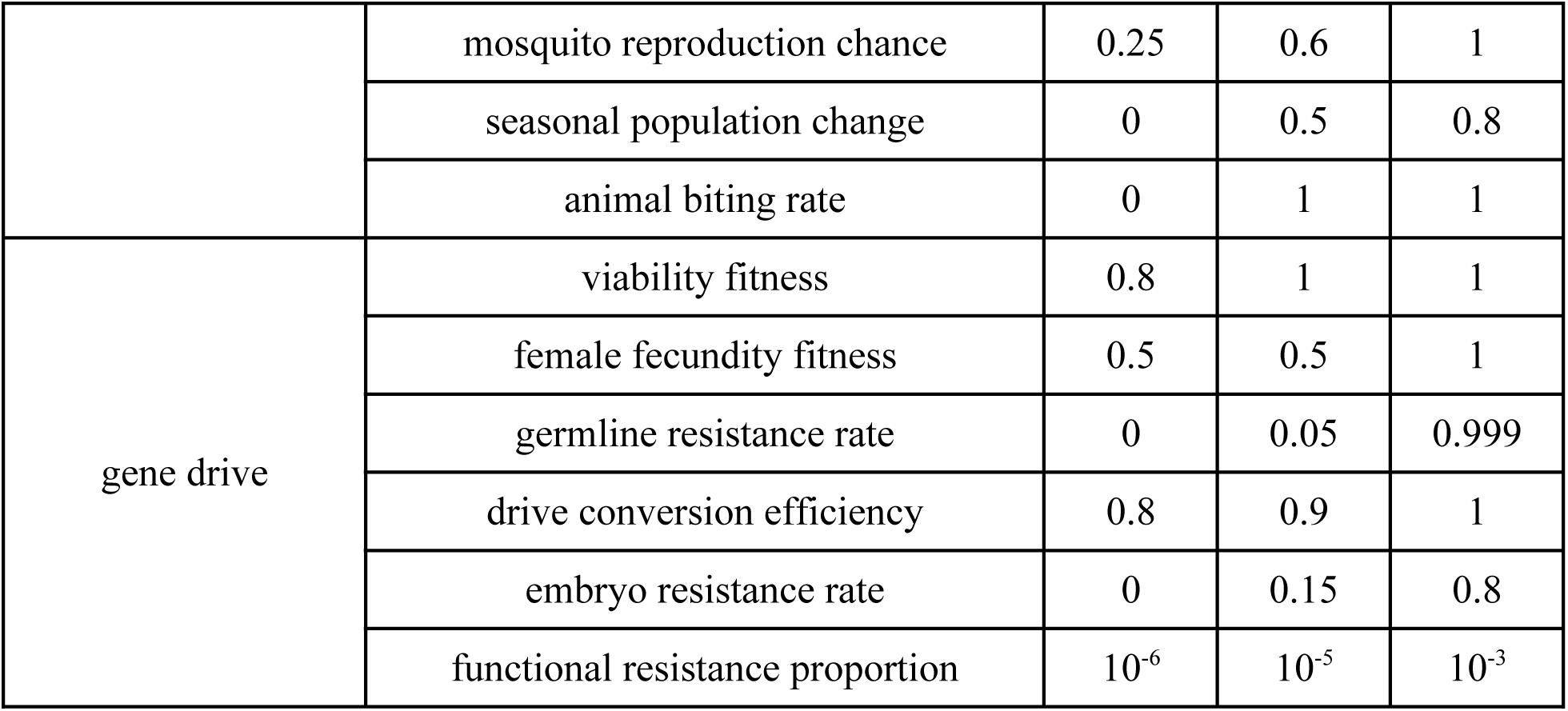
parameter range and independent default parameters.

| parameter type | parameter name | min | default | max |
| --- | --- | --- | --- | --- |
| malaria | transmission rate to mosquitoes | 0.05 | 0.11 | 0.27 |
|  | transmission rate to humans | 0.2 | 0.3 | 0.9 |
|  | infection duration (weeks) | 10 | 15 | 30 |
|  | immunity duration (weeks) | 0 | 5 | 20 |
| mosquito ecological | low-density growth rate | 2 | 6 | 18 |
|  | average dispersal(km/week) | 0.033 | 0.11 | 0.165 |
|  | weekly remating chance | 0 | 0.05 | 1 |
|  | mosquito reproduction chance | 0.25 | 0.6 | 1 |
|  | seasonal population change | 0 | 0.5 | 0.8 |
|  | animal biting rate | 0 | 1 | 1 |
| gene drive | viability fitness | 0.8 | 1 | 1 |
|  | female fecundity fitness | 0.5 | 0.5 | 1 |
|  | germline resistance rate | 0 | 0.05 | 0.999 |
|  | drive conversion efficiency | 0.8 | 0.9 | 1 |
|  | embryo resistance rate | 0 | 0.15 | 0.8 |
| | functional resistance proportion | $10^{-6}$ | $10^{-5}$ | $10^{-3}$ |

The outcomes of each simulation included whether the mosquito population and whether malaria was eliminated. Each simulation produced one vector consisting of independent parameter variables and two outcomes. By concatenating these vectors, we made a parameter and outcome matrix that could be used for our deep learning model.

We first performed 10,000 simulations to generate training and testing data for the model. To reduce the impact of a small sample size in the “success” category , we then performed an additional 10,000 simulations from an adjusted parameter space. The sample space for testing data was adjusted to restrict drive conversion to 90% or above (instead of 80%) to focus on regions of the parameter space where success was possible.

We first generated vectors of parameters not associated with gene drive as described in the next section and concatenated them with randomly generated gene drive parameters from Latin hypercube sampling, ensuring that the set of parameter points is evenly and randomly distributed in the parameter space. Because outcomes in our models were stochastic, each chosen point in the parameter space was repeated 20 times for the model without gene drive (for determining baseline malaria rates) and 10 times for the gene drive human-malaria-mosquito model. We thus obtained the probability of wildtype mosquito suppression and the probability of malaria eradication in each parameter point. The probability value was used for the regression model. For binary classification, we defined an elimination probability greater than or equal to 0.5 as “success”. We also record the malaria prevalence at week 600 and the average fertile female density during chasing. The start of chasing is defined by the first maximum point of Green’s coefficient or the first minimum of population size.

Prior to training the deep learning model, the dataset underwent preprocessing steps to ensure data quality and compatibility. We conducted normalization for all datasets. Maximum absolute scaler from Scikit-learn version 1.3.2 was used for normalization.

2.2 Preliminary analysis of human-malaria-mosquito model

We needed to ensure that the prevalence of malaria among the human population reached to an equilibrium of 30 to 50 % at week 100, representing a region with heavy malaria burden prior to the introduction of gene drive. In certain parameter sets, the simulation halted before week 100 because either mosquitoes or malaria was eradicated (thus representing unrealistic parameter sets). To address this issue, we conducted a preliminary analysis by performing the human-malaria-mosquito model simulation for 100 weeks and measuring the malaria prevalence of the last 7 weeks of the simulation. We performed SLiM simulations at 5000 sample points with 20 replicates from a Latin hypercube sampling of 10 parameters not associated with gene drive and built a deep-learning based model that estimates the average malaria prevalence from week 93 to week 100. With this model, we predicted the malaria prevalence of 100,000 random sample points and selected the points that yielded a prevalence between 30% and 50% at week 100. To improve the accuracy of training, two neural networks with different parameter sets were trained from the data simulated with the human-malaria-mosquito model. The first estimated the effect of 16 parameters on malaria prevalence, and the second estimated the effect of 12 parameters on mosquito suppression. This was to omit the effects of malaria-related parameters on the estimation of mosquito prevalence, which should have no effect.

2.3 Neural network methodology

The deep learning model was implemented with Keras version 2.12.0 on Python 3.10.11. We constructed the neural network with 16 neuron units and a number of layers varying from 1 to 5 for optimization, and we employed a linear activation function. The model was compiled with Adam optimizer using the default parameters. The model was trained for 500 epochs, where one epoch represents a complete pass through the entire dataset. The batch size was set to 32, and the validation frequency was set to 0.1. The loss function was calculated with mean squared error.

2.4 Ensemble Learning Framework

In this study, three models were trained using specific parameters for each. The LightGBM model (version 4.5.0) was configured for regression with a root mean squared error (RMSE) metric utilizing Gradient Boosting Decision Trees (GBDT). It featured a maximum of 31 leaves, a learning rate of 0.2, and a feature fraction of 90%. The XGBoost model (version 2.1.3) also focused on regression, minimizing squared error, and had a maximum tree depth of 5, a learning rate of 0.2, and a column sample of 80% for each tree. The CatBoost model (version 1.2.7) was set to perform 1000 iterations with a tree depth of 6 and a learning rate of 0.1. It provided verbose output every 100 iterations, along with an L2 leaf regularization term of 5 to mitigate overfitting. The data preprocessing methods were the same as those used for the deep learning models. For the deep learning model, twenty models were trained at each depth level from 1 to 5 layers. The results of models with the same depth were averaged to obtain the final outcomes.

2.5 Evaluation of Machine Learning Models

The performance of the model was evaluated by prediction accuracy of classification model, the coefficient of determination R^2^, Root Mean Square Error (RMSE), and Mean Average Error (MAE) for the regression model. Accuracy was evaluated with precision, recall, and F1 values, which are defined as follows:

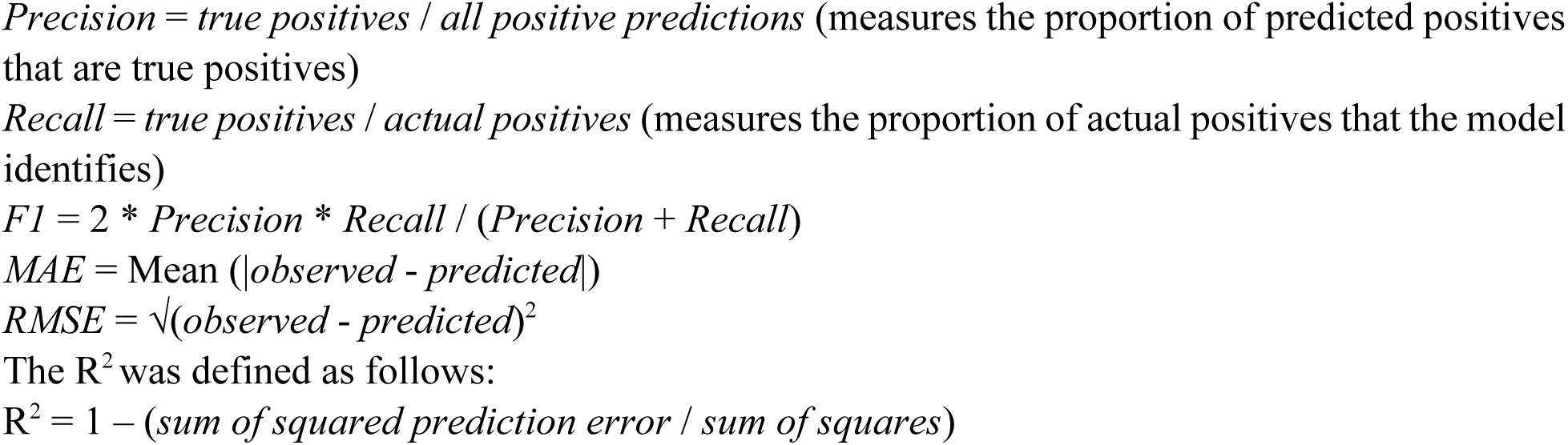

By dividing the results into regions with positive and negative classes, the model’s performance was assessed in different areas. This approach allowed for a more nuanced evaluation of how well the model predicts outcomes in regions where successful elimination occurred compared to those where it did not. The outcomes of the SLiM model are subject to randomness, and every parameter combination is represented by 10 simulations. These repeats can be regards as 10 random samples with the same expected probability. Even a perfect machine learning model that gives predictions exactly equal to the expected probabilities can’t get 100% accuracy on a test dataset with such random sampling error. To measure the gap between our machine learning model and this ideal “perfect model”, we conducted a two-sided binominal test using the predicted probability of elimination as the mean value and the observed frequency as the sample. We calculated the frequency of parameter combinations that have observed data located in the 95% confidence interval of the predicted value. The average of the residuals was used to reflect the model’s bias in predictions for both the overall dataset and the positive class of parameter combinations (cases with at least one successful elimination).

### 2.6 Estimating the effects of variables on the gene drive performance

Deep-learning is sometimes called a “black-box” — it returns a model that predicts the outcome but is difficult to interpret(Alain & Bengio, 2016). Using SHAP, a concept came from game theory, we can quantify the contribution that each variable makes to the outcome prediction(Lundberg & Lee, 2017). SHAP values show how much each parameter contributes to the prediction compared to the average prediction. With SHAP value, we can estimate which features play the most important role on the outcome. This let us better understand the logic behind the predictions of deep-learning based models. SHAP values for each variable parameter were calculated with SHAP version 0.41.0(Lundberg & Lee, 2017).

We also generated heatmaps with several combinations of two variable parameters while fixing all other values to their default levels, showing the predicted outcome values with the deep-learning based model. The default values were selected to reflect the data from previous studies, including laboratory data and ecological estimates in some cases. The heatmaps were generated with seaborn package version 0.12.2(Waskom, 2021). For malaria transmission parameters that have no influence on mosquitoes, only heatmaps showing malaria outcomes are shown. Heatmaps were generated by using the average of twenty deep learning models, each with a three-layer structure. The malaria prevalence before gene drive release is predicted by another deep learning model, which was used for selecting the training parameter range to ensure that the starting malaria proportion was within the desired range. The malaria reduction proportions are calculated based on the malaria prevalence before gene drive release.

### 3. Quantification of chasing

To investigate the potential impact of the chasing phenomenon on reduction of malaria by gene drive, we define a Chasing Index as *1−NNI*, where *NNI* is the ratio of the average nearest neighbor distance of individuals to the expected value based on a random distribution of the population(Zhang et al., 2024). A larger Chasing Index value indicates more intense chasing. To investigate the potential impact of the chasing phenomenon on malaria suppression, we collected a new dataset that simulates conditions of intense malaria transmission, incorporating several key parameters. Here, we vary several gene drive parameters and the population’s low-density growth rate with the following ranges: female fecundity fitness from 0.6 to 1, drive conversion efficiency from 0.7 to 1, embryo resistance rate from 0 to 1, germline resistance rate from 0 to 0.15, and low-density growth rate from 2 to 12. The remaining parameters were set as follows: functional resistance proportion at 0, malaria transmission rate to humans at 0.4, transmission rate to mosquitoes at 0.4, transmission rate to humans at 0.5, infection duration at 15 weeks, immunity duration at 5 weeks, average dispersal of 0.12 kilometers per week, weekly remating chance at 0.05, mosquito reproduction chance at 0.8, seasonal population change at 0, animal biting rate at 0.95, and viability fitness at 0.98. The simulations were run for 800 weeks, with the gene drive being released in week 11. All other settings remain consistent with those used in the machine learning dataset.

### 4. Data generation

All SLiM simulations were conducted using the High-Performance Computing Platform of the Center for Life Science at Peking University. Python was used for analyzing data and preparing the figures. All models and data can be accessed on GitHub (https://github.com/jchamper/Malaria-Drive-Model).

## Results

### 1. Malaria outcomes

Our simulations outcomes can be classified in several ways. First, we can have mosquito elimination or persistence, the former of which will always result in malaria elimination. If mosquitoes are not eliminated, we can still have malaria elimination or persistence. As we previously studied(J. Champer et al., 2021; Champer et al., 2022; Zhang et al., 2024), mosquito outcomes can involve the chasing phenomenon, where wild-type individuals recolonize empty space, with the drive following and causing local suppression, resulting in extinction-recolonization cycles. An alternative to chasing for less powerful drives is equilibrium, where the population is reduced, but remains at a lower level over the entire arena. We also allowed functional resistance allele formation in our simulations, which will quickly restore the population to equilibrium. In this study, we focus on ultimate mosquito population and malaria rate outcomes as the most important output of our model. In an example simulation, the wild-type population size and the malaria frequency in humans fluctuate seasonally, but then both decline sharply after release of the drive (Figure 2A). However, functional resistance alleles quickly form, and both the mosquito population and malaria infection frequency recovers. In another example, the drive reaches moderate frequency but is unable to completely eliminate the mosquito population (Figure 2B). Functional resistance alleles eventually form, but by this time, malaria has been eliminated from the simulation.

**Figure 2.**
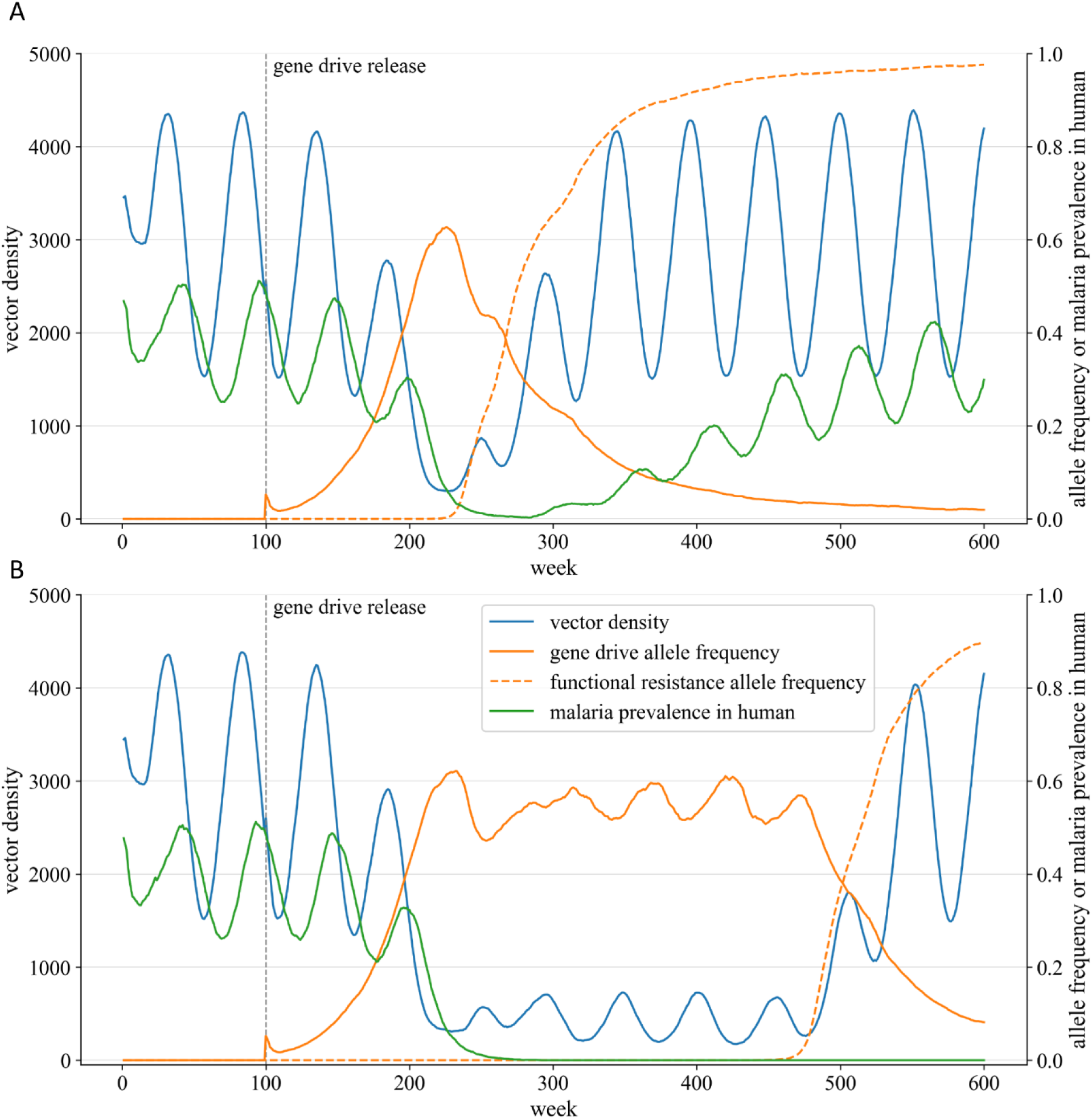
Example outcomes for gene drive and malaria. Example scenarios showing mosquito population density, malaria prevalence in humans, drive allele frequency, and resistance allele frequency over time. **A** Functional resistance allele formation quickly eliminates the drive, allowing the malaria rate to increase toward its original level. **B** The drive fails to eliminate the population, but keeps it suppressed long enough for malaria to be eliminated. Eventually, functional resistance allele formation still restores the mosquito population size.

### 2. Outcome results in model data

To obtain both the training and testing datasets for the main model, we simulated sample points that were predicted to have a malaria prevalence from 0.3 to 0.5 at week 100, estimated from the malaria prevalence model. This was to specifically simulate regions with high malaria burden. All sample points had ten replicates.

We obtained 9775 sample points in the training dataset. 143 (1.5%) of them had mosquito elimination in at least half of replicates, and 9190 (94%) of them has no replicates with mosquito elimination. 1437 (15%) parameter sets had malaria elimination in over half of replicates, and 6483 (66%) of them has no replicates with malaria elimination.

The test set contained 9776 sample points. 154 (1.6%) of them had mosquito elimination in at least half of replicates, and 9170 (94%) of them has no replicates with mosquito elimination. 1450 (15%) parameter sets had malaria elimination in over half of replicates, and 6332 (65%) of them has no replicates with malaria elimination.

### 3. Deep learning model performance

To create a deep learning model, allowing understanding of drive outcomes, we first optimized the number of hidden layers of a neural network model. We trained the model with 1 to 5 hidden 16-unit layers. We trained 20 model replicates for each type and used the average output of these models to predict results. We evaluated each model based on precision, recall, F1 value, and R^2^ value for mosquito and malaria rate suppression. Precision measures the rate of positive (elimination in at least 50% of replicates) predictions that are truly positive, while recall measures the frequency of actual positives predicted by the model. F1 is a composite model performance score based on these.

One important aspect of our data structure is that with 10 replicates per spot, our testing (and training) data is subject to stochastic variation in results. This prevents 100% model accuracy even if the model was perfect. We thus also assessed model prediction accuracy for mosquito and malaria elimination based on how many testing points fell within the 95% confidence interval of our testing data set, considering points with any mosquito/malaria elimination separately from points with zero replicates where mosquitoes or malaria was eliminated.

For mosquito elimination, precision remained good in all models, but recall was generally poor (Table 2). This is likely due to the small rate of mosquito elimination combined with stochastic outcome variation across the entire parameter set where mosquito elimination was possible. When assessing predictions using confidence intervals, thus better taking stochastic outcome variation into account, we found accuracy rates of over 80% in all models for samples with any mosquito elimination outcomes. The 3-layer deep learning model was particularly accurate, with 91% of predictions falling within testing data confidence intervals. Model accuracy was nearly 100% in regions where mosquito elimination did not occur in testing data. When elimination failed, we predicted the relative fertile adult female mosquito density (compared to initial population size) after the drive caused the first local minimum in the mosquito population. All models had R^2^ values that were usually over 0.8 (and thus fairly good, considering the stochastic variation from functional resistance allele formation, which could restore the population at a variable time). The frequency distribution and residual plots of predicted and observed values show that the deep learning model tends to somewhat underestimate the success rate in areas with high success rates in mosquito elimination (Figure S2, Figure S3).

**Table 2.** Model performance for mosquito suppression.

| model type | mosquito elimination$\geq 0.5$ | | | total | elimination $>0$ \| $=0$ | | | mosquito density | | |
| --- | --- | --- | --- | --- | --- | --- | --- | --- | --- | --- |
|  | Precision | Recall | F1 value | mean residual | mean residual | in 95% CI* | in 95% CI* | Root Mean Square Error | Mean Absolute Error | R<sup>2</sup> |
| lightBGM | 0.831 | 0.318 | 0.460 | -0.003 | 0.082 | 0.823 | 0.998 | 0.064 | 0.027 | 0.822 |
| XGBoost | 0.787 | 0.240 | 0.368 | -0.003 | 0.097 | 0.814 | 0.999 | 0.065 | 0.029 | 0.817 |
| CatBoost | 0.857 | 0.351 | 0.498 | -0.003 | 0.076 | 0.880 | 0.999 | 0.060 | 0.026 | 0.842 |
| DL 1-layer | 0.857 | 0.312 | 0.457 | -0.001 | 0.090 | 0.828 | 0.999 | 0.072 | 0.034 | 0.776 |
| DL 2-layer | 0.867 | 0.506 | 0.639 | -0.002 | 0.053 | 0.888 | 1.000 | 0.063 | 0.025 | 0.829 |
| DL 3-layer | 0.874 | 0.584 | 0.700 | -0.003 | 0.049 | 0.911 | 1.000 | 0.060 | 0.025 | 0.843 |
| DL 4-layer | 0.878 | 0.513 | 0.648 | 0.001 | 0.078 | 0.866 | 1.000 | 0.060 | 0.023 | 0.845 |
| DL 5-layer | 0.861 | 0.565 | 0.682 | 0.000 | 0.061 | 0.884 | 1.000 | 0.059 | 0.025 | 0.846 |

\*The fraction of observed data located in the 95% confidence interval of the predicted elimination rate.

We conducted a similar analysis on malaria elimination and prevalence (Table 3). For these models, precision and recall was substantially improved, likely due to smaller stochastic variation in malaria elimination outcomes. Prediction of elimination rates for the testing parameter space with any elimination was moderately improved compared to the mosquito elimination model (with the 3-layer deep learning model again having the best performance of 94%), though predictions in parameter space no elimination were slightly worse (though still accurate) compared to mosquito elimination, perhaps due to small numbers of success in parameter ranges where the drive would consistently fail due to functional resistance or chasing, but still retain the ability to stochastically eliminate malaria in some cases. Malaria reduction prediction R^2^ values were also somewhat higher than for mosquito suppression. In terms of distribution, the 3-layer deep learning model had little deviation in predicting malaria elimination rate across the range (Figure S4, Figure S5).

**Table 3.** Model performance for malaria prevalence.

| model type | malaria elimination$\geq 0.5$ | | | total | elimination $>0$ \| $=0$ | | | malaria prevalence | | |
| --- | --- | --- | --- | --- | --- | --- | --- | --- | --- | --- |
|  | Precision | Recall | F1 value | mean residual | mean residual | in 95% CI* | in 95% CI* | Root Mean Square Error | Mean Absolute Error | R<sup>2</sup> |
| lightBGM | 0.911 | 0.717 | 0.803 | -0.004 | 0.043 | 0.892 | 0.991 | 0.062 | 0.047 | 0.828 |
| XGBoost | 0.923 | 0.694 | 0.793 | -0.005 | 0.045 | 0.911 | 0.989 | 0.067 | 0.052 | 0.802 |
| CatBoost | 0.931 | 0.718 | 0.811 | -0.005 | 0.038 | 0.919 | 0.993 | 0.049 | 0.035 | 0.894 |
| DL 1-layer | 0.942 | 0.686 | 0.794 | -0.001 | 0.050 | 0.919 | 0.994 | 0.064 | 0.049 | 0.818 |
| DL 2-layer | 0.920 | 0.773 | 0.840 | -0.001 | 0.032 | 0.932 | 0.994 | 0.055 | 0.041 | 0.864 |
| DL 3-layer | 0.910 | 0.786 | 0.844 | -0.003 | 0.027 | 0.941 | 0.995 | 0.053 | 0.039 | 0.876 |
| DL 4-layer | 0.924 | 0.767 | 0.838 | 0.003 | 0.038 | 0.933 | 0.996 | 0.053 | 0.040 | 0.877 |
| DL 5-layer | 0.920 | 0.782 | 0.846 | 0.001 | 0.034 | 0.938 | 0.995 | 0.054 | 0.040 | 0.874 |

\*The fraction of observed data located in the 95% confidence interval of the predicted elimination rate.

### 4. Effect of specific parameters on mosquito and malaria suppression

Because the 3-layer deep learning model appeared to be the most accurate across most measures for both mosquito and malaria suppression, we selected it for further analysis. We first predicted the effect of varying pairs of parameters while keeping others at their default values.

The drive conversion rate is the most important parameter of gene drive performance. Even when restricting it to high values (80%+), mosquito elimination was rare (Figure 3), though substantial population reduction could still be obtained in a ten-year interval despite possible formation of functional resistance alleles. Suppression drives often have reductions in female fitness due to somatic expression or other disruption of the target gene in required tissues. This could also severely degrade performance, if to a lesser extent than drive efficiency. However, over our selected parameter range, malaria elimination was nearly certain for drives with optimal performance and was still likely even with drive efficiency and female fitness at the low end of our parameter range (Figure 3).

**Figure 3.**
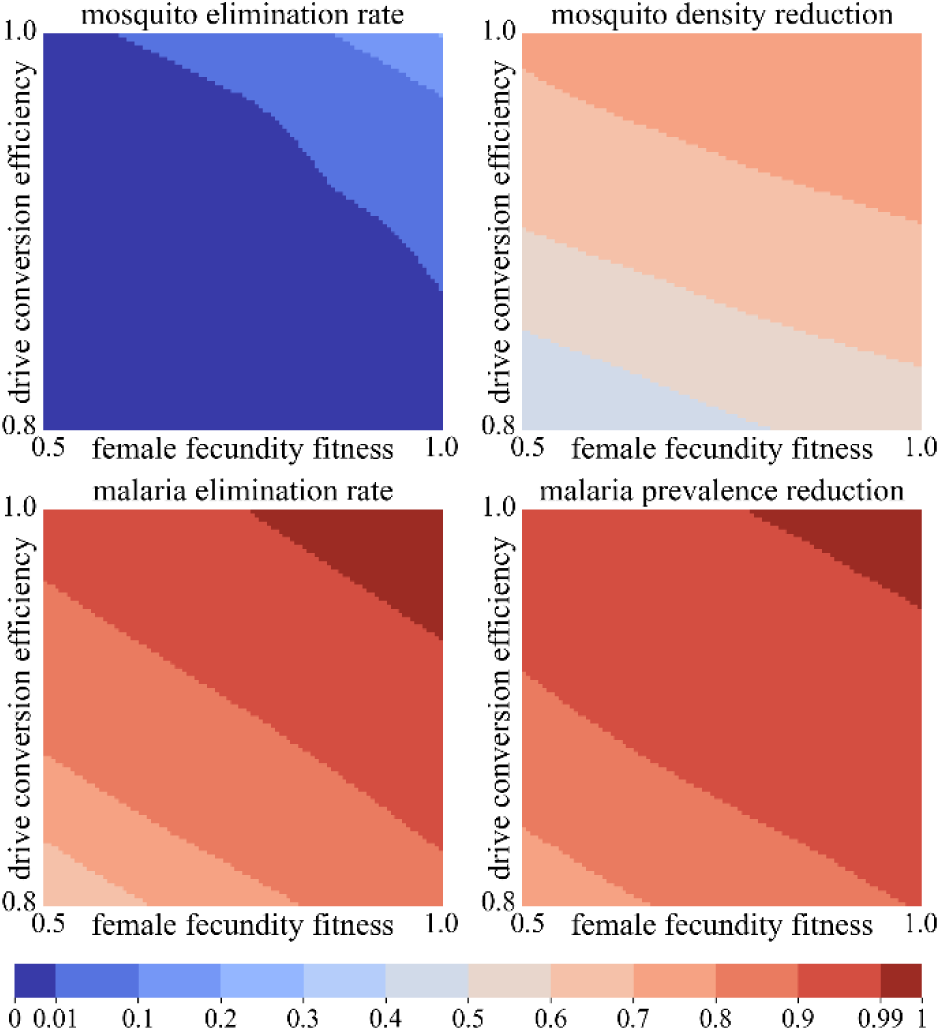
Effect of drive performance parameters. Gene drive outcomes were predicted using the 3-layer deep learning model. Parameters are fixed at their default varies, with varying drive conversion efficiency and fecundity fitness of female drive heterozygotes. We show the mosquito elimination rate, mosquito density reduction, malaria elimination rate, and malaria prevalence reduction compared to the pre-drive average.

Even if a drive was potentially capable of eliminating malaria, resistance could prevent it. Nonfunctional resistance can allow a higher mosquito population at equilibrium, and functional resistance could quickly result in drive failure and revert the population back to its original size. While the latest *Anopheles* drives have little embryo resistance (under 20%), this still represents the primary source of resistance alleles(Carballar-Lejarazú et al., 2022). At our default parameters, embryo resistance would only have a major effect at higher values (Figure 4). However, if an intermediate proportion (over 10^-5^ to 10^-4^) of resistance alleles result in a functional target gene, both mosquito and malaria reduction become very unlikely. For populations larger than the one that we modeled, the fraction of functional resistance alleles may need to be proportionally lower to avoid similar drive failure outcomes.

**Figure 4.**
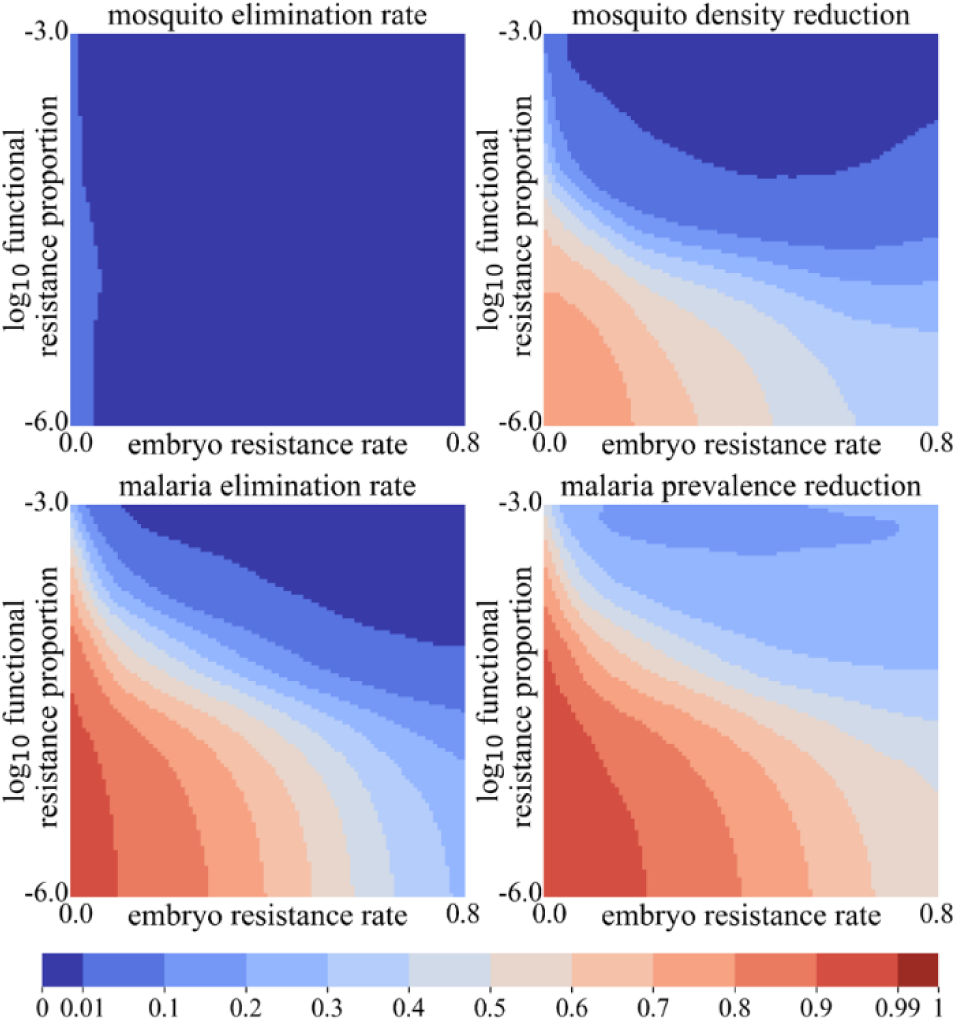
Effect of resistance alleles. Gene drive outcomes were predicted using the 3-layer deep learning model. Parameters are fixed at their default varies, with varying embryo resistance allele formation rate and relative proportion of functional resistance alleles. We show the mosquito elimination rate, mosquito density reduction, malaria elimination rate, and malaria prevalence reduction compared to the pre-drive average.

Previous studies on chasing indicated that it could often be avoided if the average dispersal rate was sufficiently high(J. Champer et al., 2021; Champer et al., 2022). However, when holding other parameters at their default, high dispersal still did not allow much greater success in our simulations, having only a modest effect in reducing the average fertile female population size at higher values (Figure 5). However, when dispersal was low, malaria elimination was high. This is partially because malaria elimination become substantially slowed at low dispersal, and indeed very low values were outside our data set parameter range because they produced a malaria prevalence of below 30%. The low-density growth rate had a somewhat larger effect, as expected because higher values tend to produce a greater number of mosquito females during drive-induced chasing or equilibrium (Figure 5). It is unclear exactly what the low-density growth rate should be for *Anopheles* models. Higher values may best represent laboratory populations, but in the field, competing species and predators may substantially reduce the effective low density growth rate for purposes of suppression modeling(Liu et al., 2023). If it becomes low enough, the chance of malaria and mosquito elimination are notably improved.

**Figure 5.**
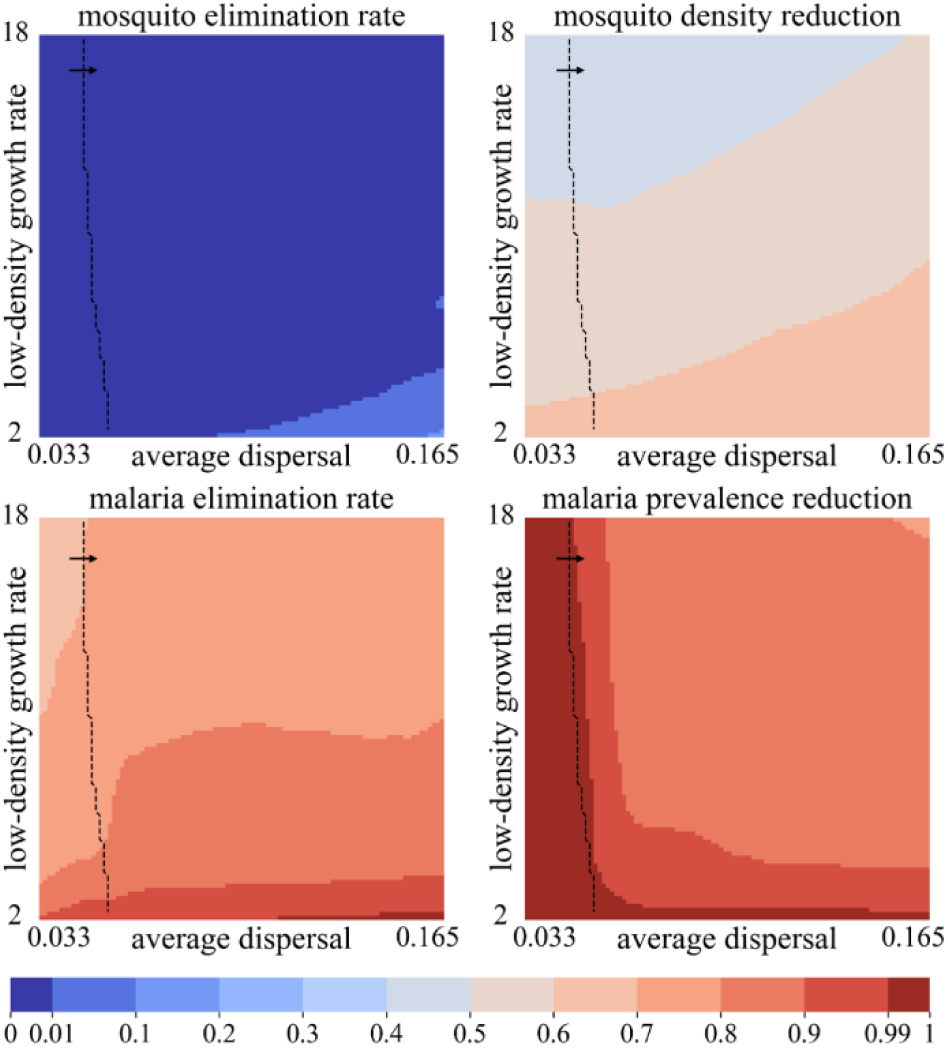
Effect of mosquito ecology parameters. Gene drive outcomes were predicted using the 3-layer deep learning model. Parameters are fixed at their default varies, with varying average dispersal (km traveled by adults per week) and low-density growth rate (the relative increase in the population per generation at very low density). We show the mosquito elimination rate, mosquito density reduction, malaria elimination rate, and malaria prevalence reduction compared to the pre-drive average. The deep learning model is trained using parameters with a malaria equilibrium prevalence in humans of between 30% to 50%. The arrow in the dashed lines shows where this parameter range is located on the heatmap (predictions outside this range may be less accurate).

Aside from these examples, our deep learning model allows rapid generation of any other pairs of parameters in our model (Figures S6–14). Together, these allow a greater understanding of the effect of individual parameters.

### 5. Contribution of model parameters for vector and disease elimination

We calculated the SHAP values for our model to better understand how individual parameters affected mosquito and malaria elimination (Figure 6). For mosquito elimination, the possible parameter range for success was small, so none of the SHAP values were high. Nonetheless, the embryo resistance allele formation rate, low-density growth rate, and drive conversion efficiency were all featured. In our simulations, these all needed to be at extreme values to have a substantial chance of mosquito elimination in the first place. Only then could other factors potentially reduce the chance of elimination, such as functional resistance and dispersal (including weekly remating, which increases the effective dispersal value of alleles).

**Figure 6.**
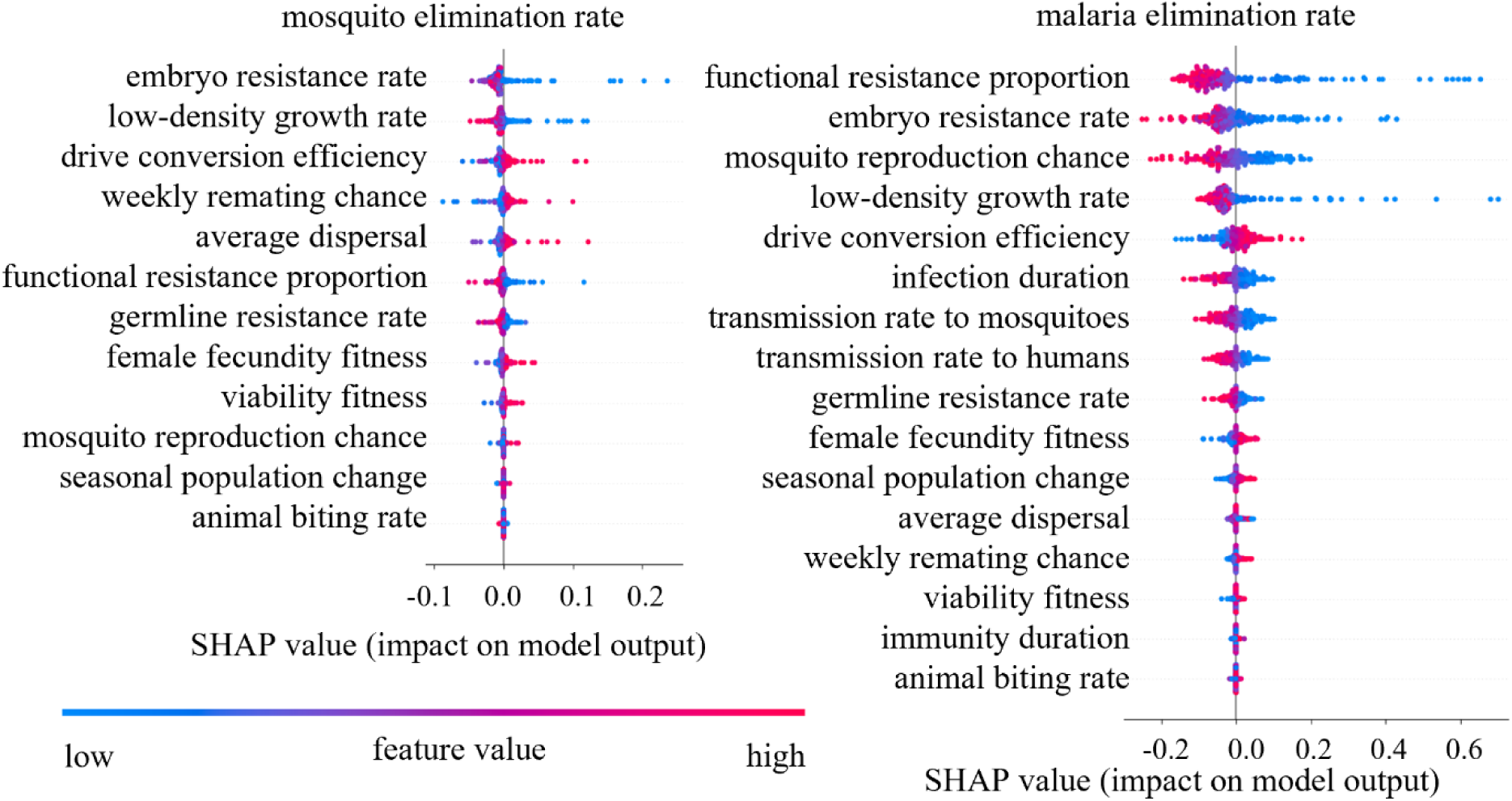
SHAP values for mosquito and malaria elimination. SHAP values for 12 parameters for mosquito elimination and 16 parameters for malaria elimination are shown. The first 200 parameter combinations are used to calculate SHAP values, and greater values indicate a positive contribution to the outcome. Red represents high feature value, and blue represents low feature value. All features are sorted by average absolute value of the SHAP value.

Malaria elimination occurred over a wider parameter range, so more parameters could have a larger magnitude of effect. Functional resistance was the most important, followed by embryo resistance (perhaps mostly from its indirect contribution to functional resistance). Also important were mosquito reproduction factors. Low-density growth rate allows their population to remain high even in the face of a strong suppression drive, and the mosquito reproduction chance allows higher biting rate, increasing malaria transmission between human and mosquitoes. Drive conversion and infection parameters were also fairly important. Many of these aspects cannot be directly controlled, but these results nonetheless further emphasize the need to reduce functional resistance allele formation. It also shows that changes in malaria parameters can aid in malaria elimination, implying that a gene drive campaign and more intensive malaria detection and treatment may have synergistic positive effects.

### 6. Effect of chasing on malaria transmission

Based on our previous results of high rates of chasing in the mosquito model(Champer et al., 2022; Zhang et al., 2024) together with moderate rates of malaria elimination in this study, we were curious about the effect of chasing on malaria prevalence. We thus conducted new simulations, representing a high-malaria region, where we measured the chasing index. This is 1 – “average nearest neighbor index,” as studied recently(Zhang et al., 2024), which represents the local non-uniformity of population distribution caused by gene drive. A high chasing index means that individuals in the population are distributed in smaller spatial clusters, so chasing is “stronger” in these simulations for the same population size. An index of zero means that individuals are almost uniformly distributed in the simulation area, representing a more stable equilibrium. As indicated by our results, the malaria prevalence is correlated with vector density (Figure 7). However, we also found that in simulations with higher chasing index, malaria prevalence is often substantially reduced or eliminated. This may indicate that even though chasing can often prevent population elimination, it may not contribute much to malaria persistence.

**Figure 7.**
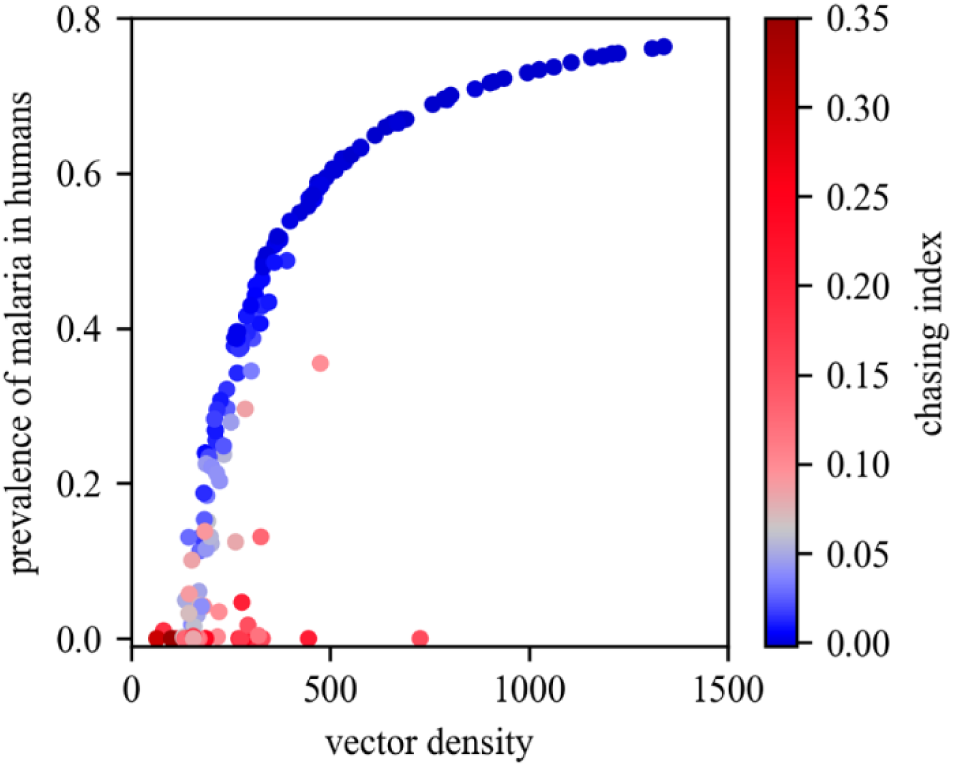
The effect of chasing on malaria elimination. Several simulations were performed with varying gene drive parameters and low-density growth rate. The average vector density (of just fertile adult females) and malaria prevalence are plotted for the last two years of the simulation. Points are colored by the chasing index (1 – average nearest neighbor index), which represents the magnitude of chasing in the simulation, with zero representing an equilibrium outcome.

## Discussion

In this study, we used a machine-learning model to assess a spatially explicit, individual-based model of homing suppression drive with malaria transmission between mosquitoes and humans. We found that mosquito elimination requires exceptionally high drive performance, but that malaria elimination was considerably easier. It was possible even if the gene drive failed due to chasing or functional resistance allele formation.

Due to the large number of parameters in our model, we were unable to symmetrically interrogate it by running sufficiently large numbers of simulations. Thus, we used a deep-learning framework based on a more limited number of simulations to predict outcomes over the entire relevant parameter space. This was complicated by the small number of outcomes resulting in successful mosquito population elimination, which may have stemmed from the relatively high rate of functional resistance allele formation rates in many of our simulations. This was overcome in a previous study involving rat suppression by utilizing a separate model that did not include functional resistance(S. E. Champer et al., 2021). However, by generating many different deep-learning models and taking their average result, we achieved relatively good accuracy. This is likely enough for more general conclusions about possible outcomes for gene drive, but for specific gene drives, narrower parameter ranges for known values will provide higher accuracy. Increased knowledge of mosquito ecology and malaria transmission could also improve model accuracy for specific regions. This faster running deep learning model also has the potential to help slower running simulation models with parameter fitting, helping further improve the accuracy of the model as we obtain more accurate real-world data(Mondal et al., 2025).

Consistent with previous work, we found that successful elimination of mosquitoes was determined by two aspects of the model. Our SHAP results show that the low-density growth rate is a critical parameter, representing the required suppressive power of the drive. This can only be overcome by a drive with sufficiently high efficiency and low fitness costs, with nonfunctional resistance allele formation rates being of secondary importance. Second, a completely successful drive must also avoid functional resistance allele formation, which is determined by both the relative functional resistance rate and the overall resistance allele formation rates in both the germline and the embryo. In *Anopheles* at least, drive conversion rates tend to be very high, leaving little room for germline resistance allele formation(Adolfi et al., 2020; Carballar-Lejarazú et al., 2023; Carballar-Lejarazú et al., 2020; Hammond et al., 2016; Kyrou et al., 2018). However, embryo resistance remains a possibility. Thus, efforts are needed to reduce the relative rate of functional resistance, which can effectively be achieved with multiplexed gRNAs(Champer et al., 2020), as demonstrated recently(Morianou et al., 2024; Xu et al., 2025). This could allow for substantially higher success rates of realistically achievable real-world *A. gambiae* drives than shown in our model. Comparing to a study on modification gene drives to reduce malaria, our result emphasized that drive conversion efficiency is more important for suppression gene drives because drive power is a major limitation, while drive fitness plays more important role in modification gene drives, where the drive is more likely to contend with functional resistance after it spreads to most of the population(Mondal et al., 2024b).

Mosquito ecology remains a critical point of consideration, yet despite efforts, many basic parameters remain poorly understood. The low-density growth rate is one of these, as is the exact shape of the density growth curve. In our model, we used a conservative density growth curve, but other curves could increase the number of mosquitoes during chasing(Zhang et al., 2024). Both of these could also vary seasonally, which we did not model (only population capacity varied with season in our model). Further complicating these are interspecies ecological interactions. For example, competing species or predators could reduce the effective low-density growth rates and considerably ease population elimination by an imperfect gene drive(Liu et al., 2023). If a competing species can also serve as a vector, though, malaria elimination may be further complicated, even if target mosquito species elimination is easier.

Most malaria elimination in our model occurred in parameter ranges where the drive itself failed due to chasing, which was often followed by functional resistance allele formation. We found that even though chasing makes mosquito elimination considerably more difficult, it can actually contribute to malaria elimination, at least compared to similarly sized populations at a stable equilibrium. If populations are the same size, then a chasing population that is moving over a landscape has a much lower chance to transmit malaria to a human and then still have mosquitoes in the area to receive the malaria after it becomes transmissible. This can break the cycle of transmission and thus assist with malaria elimination. Population suppression for other vector-borne diseases may similarly benefit, though this is not applicable to population suppression scenarios where it is the target species itself causing the negative effect.

Though our model can simulate gene drive and malaria transmission in high detail, it nonetheless remains an approximation of potential real-world scenarios. Most notable is the small spatial scale, which is necessary due to computational limitations of individual-based spatial models, despite our deep-learning framework. While perhaps accurate locally, this model does not account for potential migration of mosquitoes or disease (or even drive activity) from outside the area, which may be important in outcomes, particularly if functional resistance results in only temporary malaria elimination, followed by reintroduction from the outside. Our model also does not explicitly take into account other mosquito and malaria control programs, though these could perhaps be represented by variations in disease transmission parameters. We also modeled an area with year-round mosquitoes, and though we included large population fluctuations due to seasonality, actual fluctuations in many regions are far higher (often involving complete mosquito elimination for several months, at least for adult mosquitoes), which likely has a large effect on both mosquito and disease transmission dynamics. Yet, our model also has unique advantages, The individual-based continuous space framework that produces a high computational burden also potentially allows a more accurate simulation of many real-world areas where mosquitoes are distributed roughly continuously over several kilometers of terrain.

Overall, our results indicate that suppression gene drives remain quite promising for malaria control. Surprisingly, even substantially less efficient systems may still carry enough power to eliminate malaria, at least in a local region. More detailed models with larger spatial scales are needed to further confirm this result and check its stability, particularly if functional resistance forms in a region. Future studies may find that our conclusions can even hold in other species and for elimination of other vector-borne diseases with imperfect drives.

## Acknowledgements

This study was supported by the Center for Life Sciences and the National Natural Science Foundation of China (32270672, W2432018).

## Supplemental Information

**Table S1.** Model constants.

| parameter type | parameter name | value | unit |
| --- | --- | --- | --- |
| malaria | latent period in human | 2 | week |
| mosquito ecological | female lifespan | 7 | week |
|  | male lifespan | 5 | week |
|  | simulation area side length | 4 | kilometer |
|  | human density | 300 | individual/km<sup>-2</sup> |
|  | competition distance | 0.033 | km |
|  | adult female density | 3000 | individual/km<sup>-2</sup> |
|  | old larva competition coefficient | 5 | - |
|  | maximum offspring number | 36 | individual |
| gene drive | release ratio | 0.01 | - |

**Figure S1.**
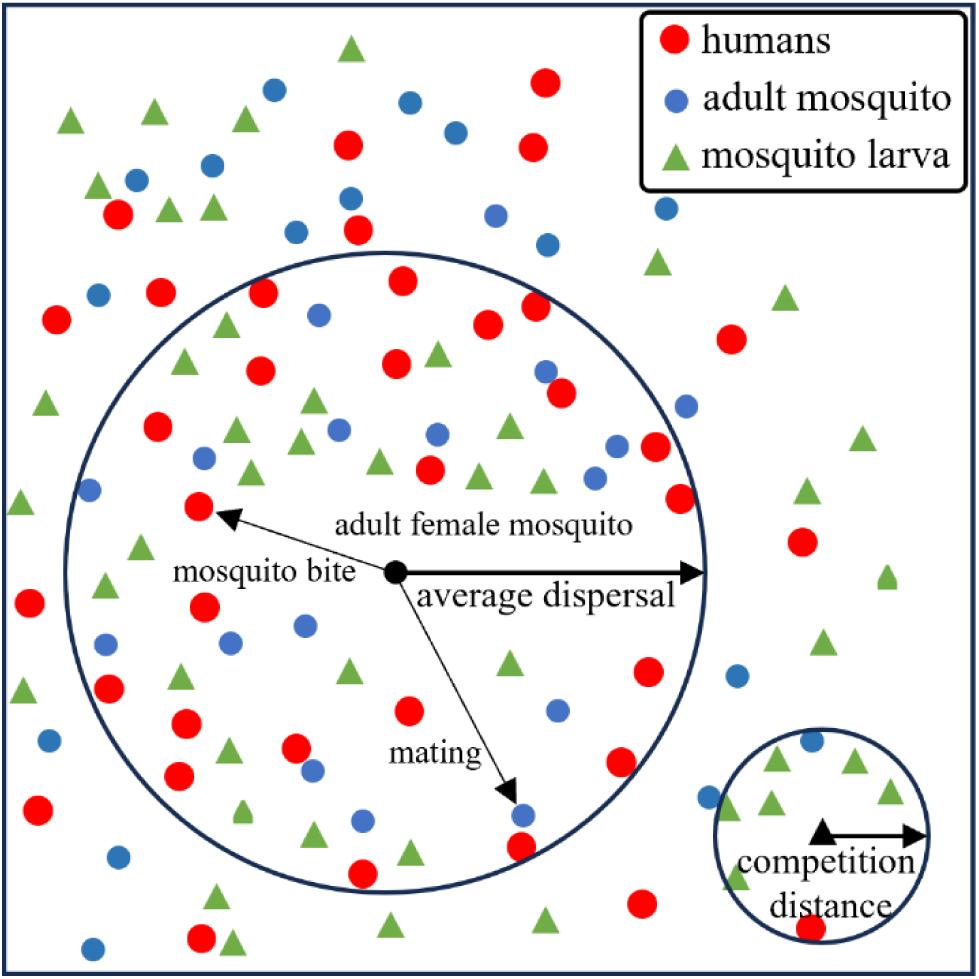
Spatial Model diagram. In each reproduction cycle, if a fertile female mosquito hasn’t mated or decided to mate again, it will mate with a random adult male mosquito around it. Then it will bite a random human(host) around it. The maximum biting and mating radius is set to the average dispersal distance (0.11 km as default). Larva competition also happens locally, with a much smaller competition distance (0.033 km).

**Figure S2.**
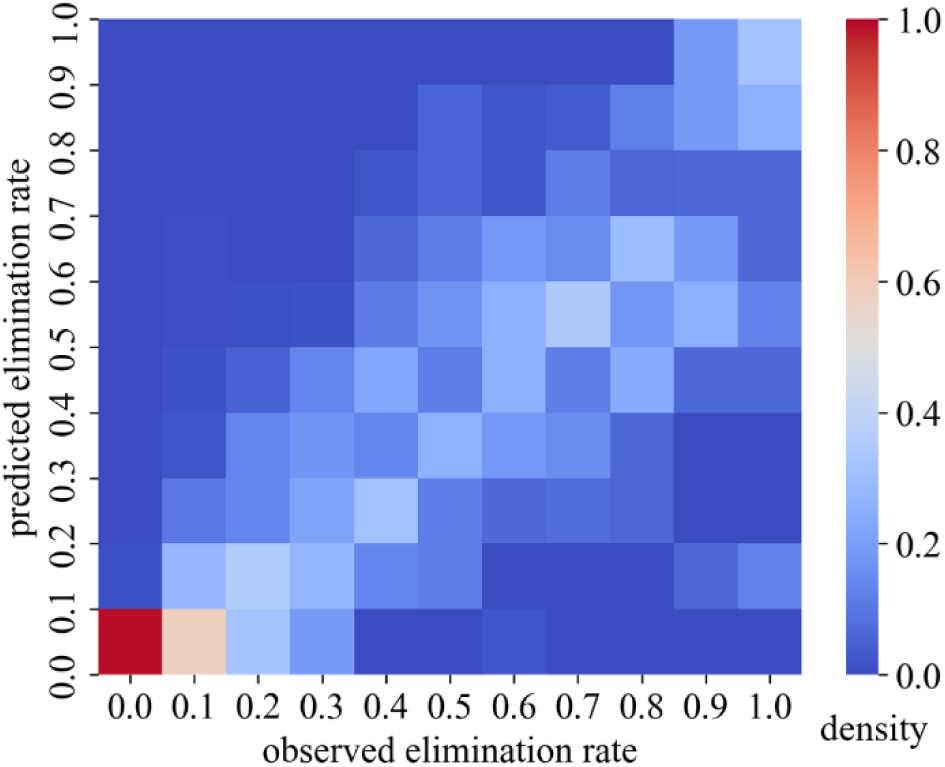
The frequency distribution of predicted mosquito elimination rate under the corresponding interval. Distribution density plot of predicted and observed values for mosquito elimination rates from the machine learning model, which is the average of twenty three-layer deep learning models.

**Figure S3.**
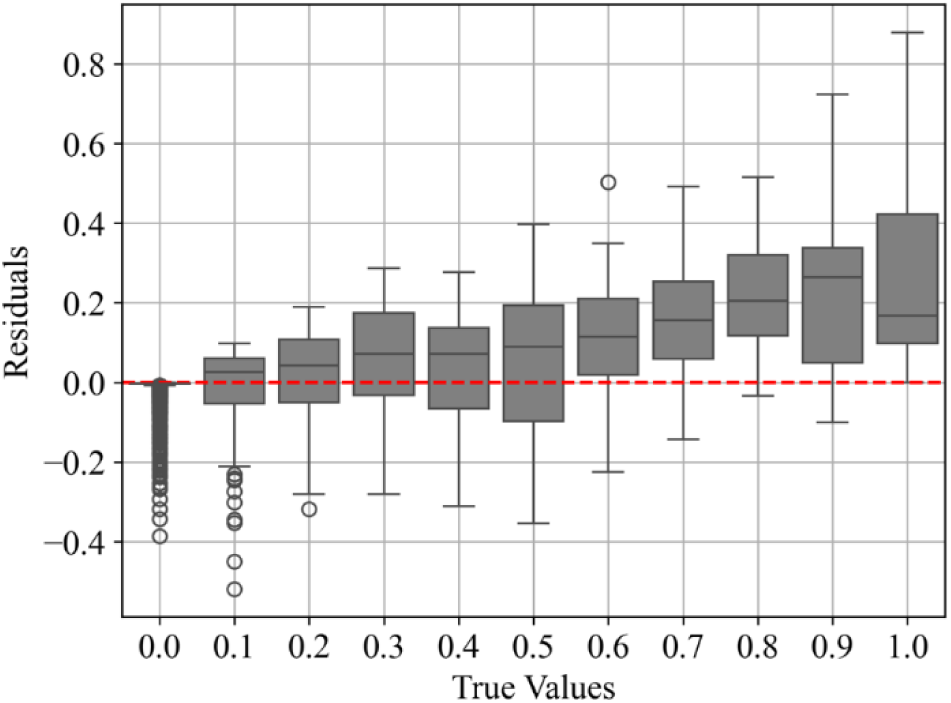
Residuals in the mosquito elimination model. Plot of residuals for the 3-layer deep learning mosquito elimination model.

**Figure S4.**
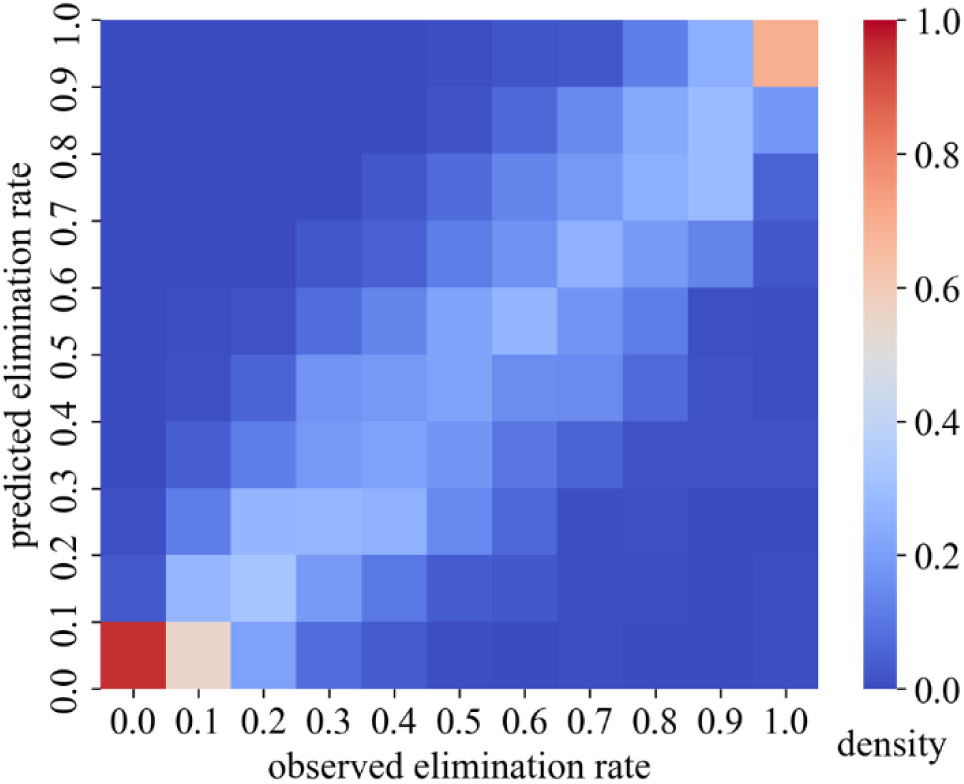
The frequency distribution of predicted malaria elimination rate under the corresponding interval. Distribution density plot of predicted and observed values for malaria elimination rates from the machine learning model, which is the average of twenty three-layer deep learning models.

**Figure S5.**
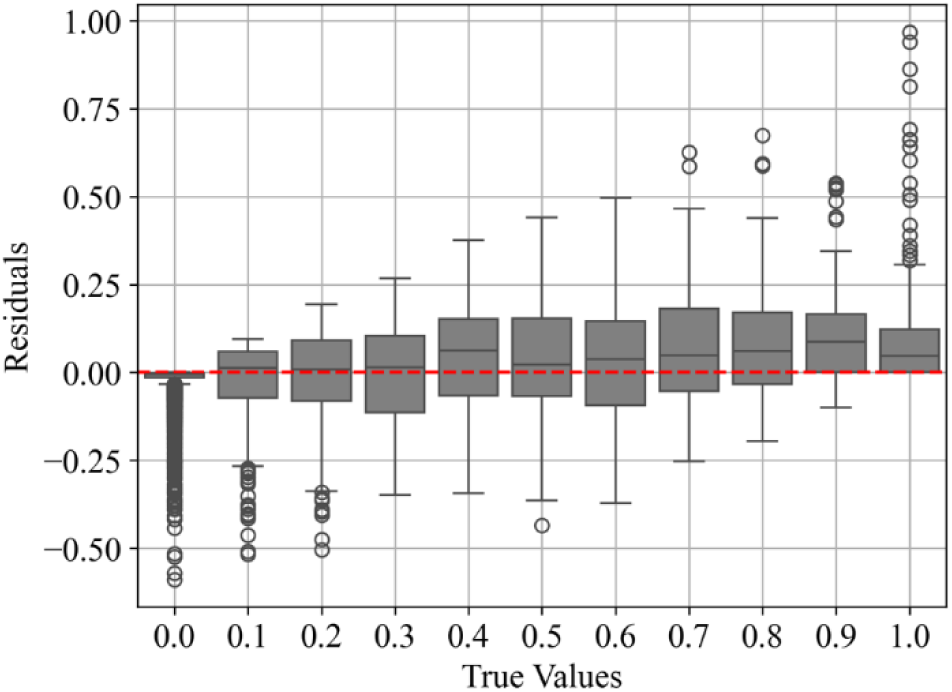
Residuals in the malaria elimination model. Plot of residuals for the 3-layer deep learning mosquito elimination model.

**Figure S6.**
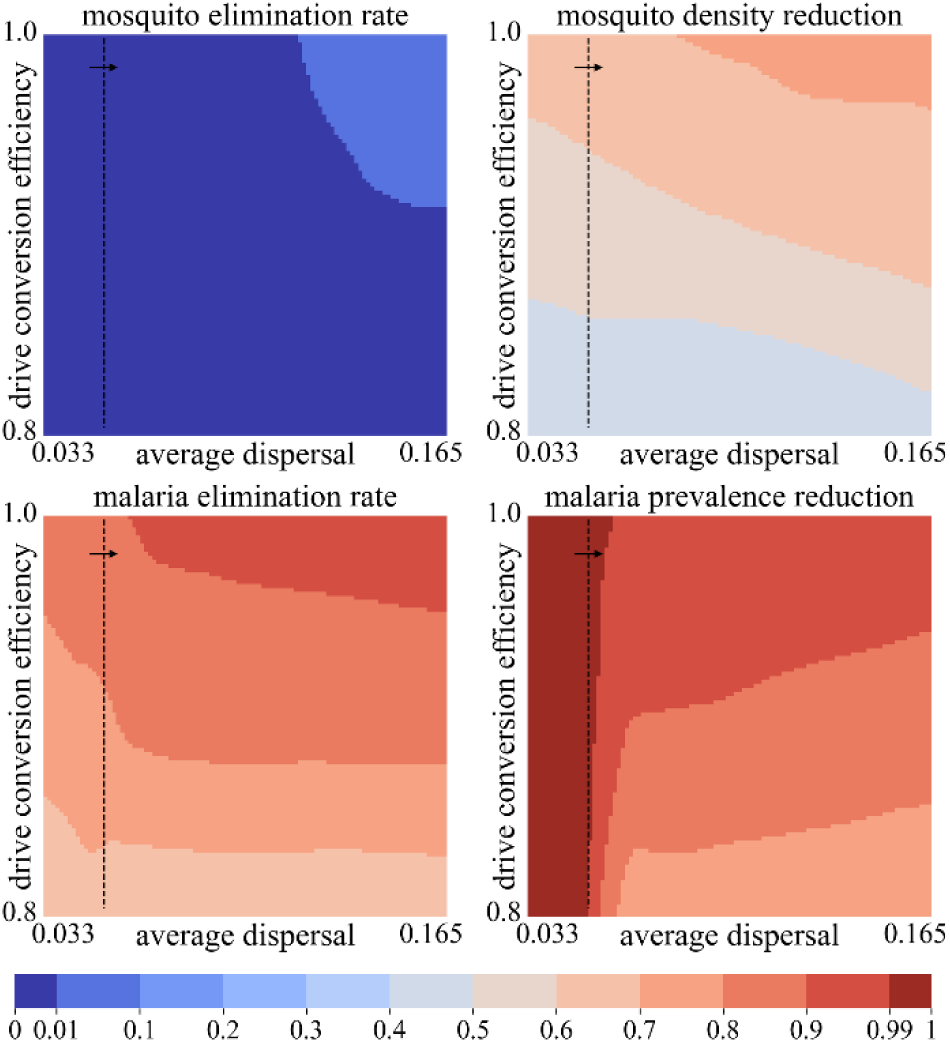
Effect of dispersal and drive efficiency. Gene drive outcomes were predicted using the 3-layer deep learning model. Parameters are fixed at their default varies, with varying average dispersal (km traveled by adults per week) and drive conversion efficiency. We show the mosquito elimination rate, mosquito density reduction, malaria elimination rate, and malaria prevalence reduction compared to the pre-drive average. The deep learning model is trained using parameters with a malaria equilibrium prevalence in humans of between 30% to 50%. The arrow in the dashed lines shows where this parameter range is located on the heatmap (predictions outside this range may be less accurate).

**Figure S7.**
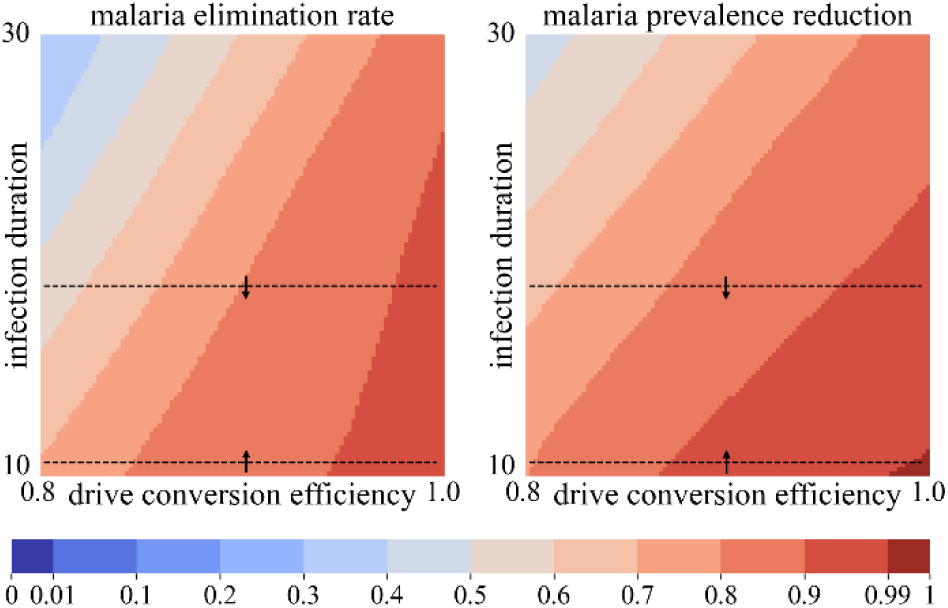
Effect of drive efficiency and infection duration. Gene drive outcomes were predicted using the 3-layer deep learning model. Parameters are fixed at their default varies, with varying drive conversion efficiency and average infection duration (weeks, of malaria in humans). We show the malaria elimination rate and malaria prevalence reduction compared to the pre-drive average. The deep learning model is trained using parameters with a malaria equilibrium prevalence in humans of between 30% to 50%. The arrow in the dashed lines shows where this parameter range is located on the heatmap (predictions outside this range may be less accurate).

**Figure S8.**
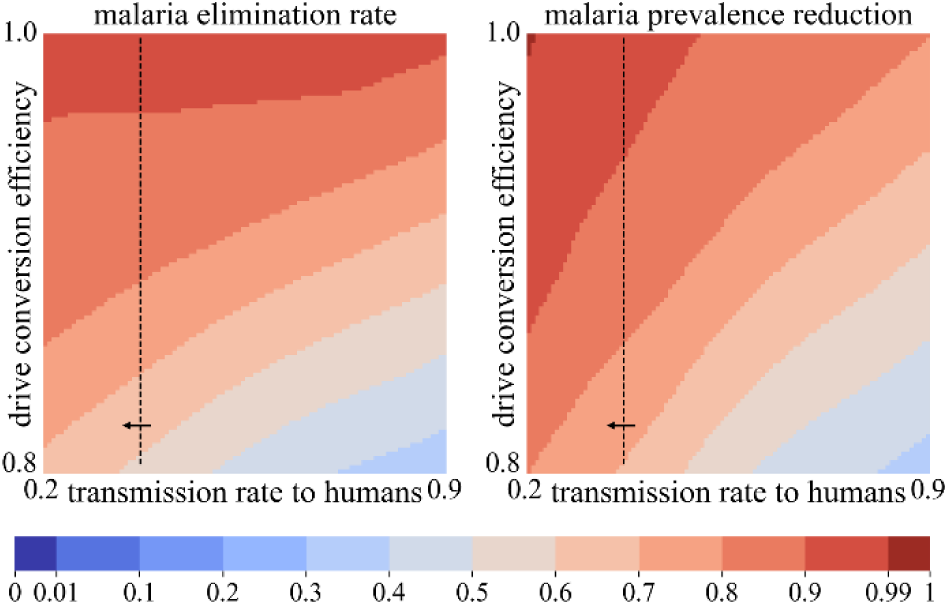
Effect of transmission rate and drive efficiency. Gene drive outcomes were predicted using the 3-layer deep learning model. Parameters are fixed at their default varies, with varying malaria transmission rate from mosquitoes to humans per bite and varying drive conversion efficiency. We show the malaria elimination rate and malaria prevalence reduction compared to the pre-drive average. The deep learning model is trained using parameters with a malaria equilibrium prevalence in humans of between 30% to 50%. The arrow in the dashed lines shows where this parameter range is located on the heatmap (predictions outside this range may be less accurate).

**Figure S9.**
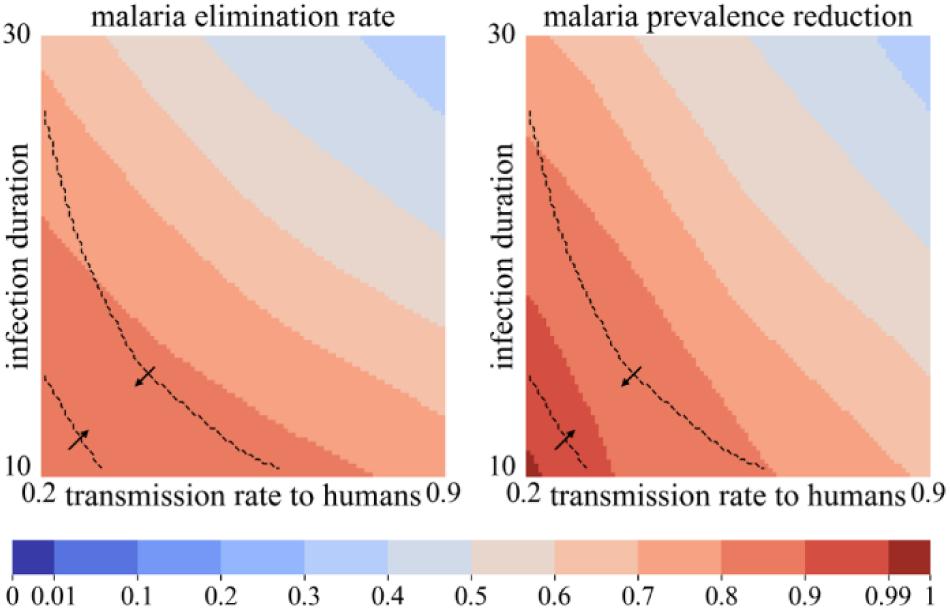
Effect of transmission rate and infection duration. Gene drive outcomes were predicted using the 3-layer deep learning model. Parameters are fixed at their default varies, with varying malaria transmission rate from mosquitoes to humans per bite and varying average infection duration (weeks, of malaria in humans). We show the malaria elimination rate and malaria prevalence reduction compared to the pre-drive average. The deep learning model is trained using parameters with a malaria equilibrium prevalence in humans of between 30% to 50%. The arrow in the dashed lines shows where this parameter range is located on the heatmap (predictions outside this range may be less accurate).

**Figure S10.**
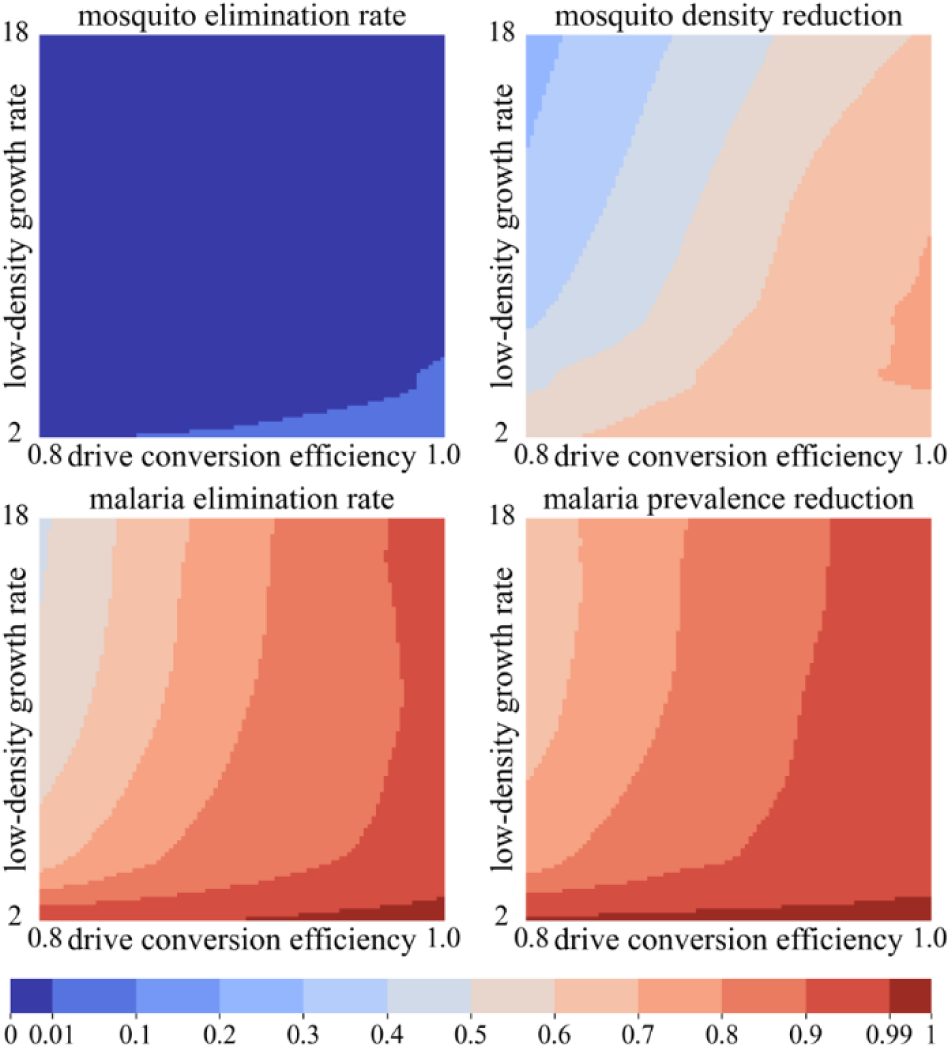
Effect of drive efficiency and low-density growth rate. Gene drive outcomes were predicted using the 3-layer deep learning model. Parameters are fixed at their default varies, with varying drive conversion efficiency and low-density growth rate. We show the mosquito elimination rate, mosquito density reduction, malaria elimination rate, and malaria prevalence reduction compared to the pre-drive average.

**Figure S11.**
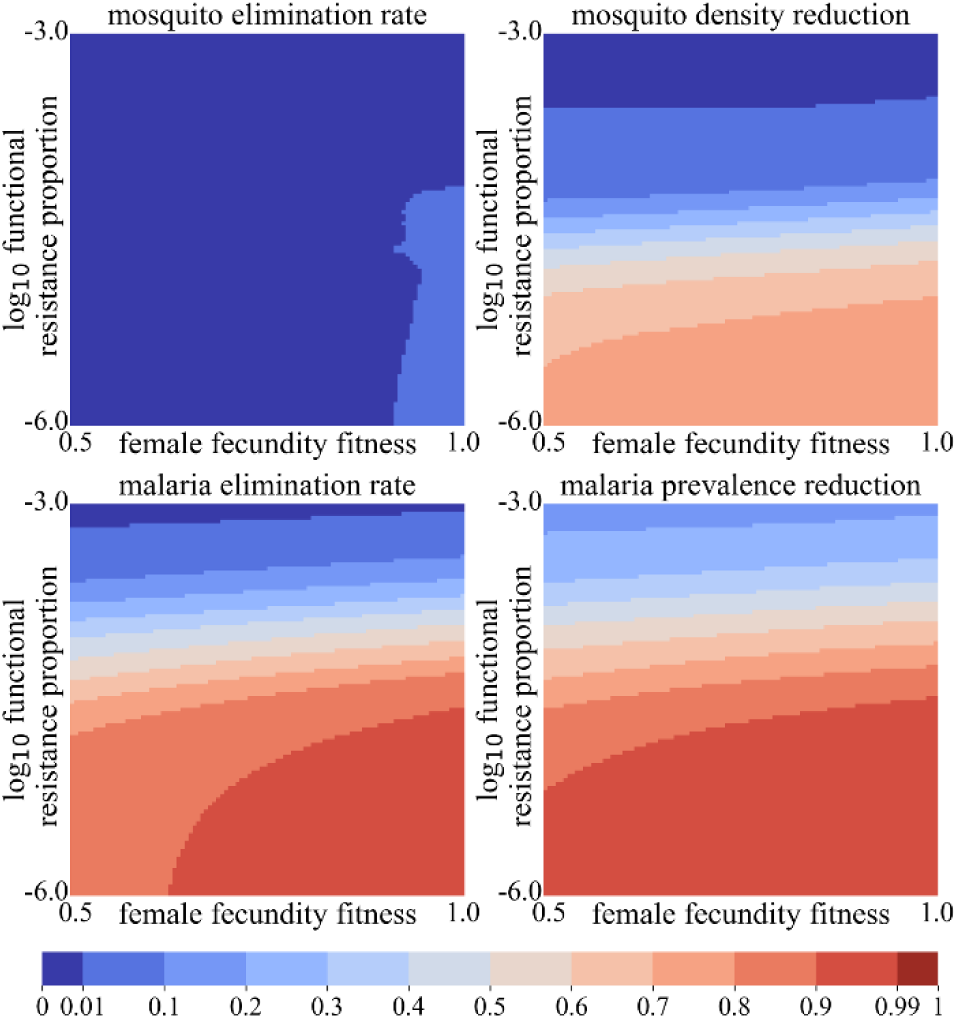
Effect of female heterozygote fitness and functional resistance. Gene drive outcomes were predicted using the 3-layer deep learning model. Parameters are fixed at their default varies, with varying fecundity fitness of female drive heterozygotes and relative proportion of functional resistance alleles. We show the mosquito elimination rate, mosquito density reduction, malaria elimination rate, and malaria prevalence reduction compared to the pre-drive average.

**Figure S12.**
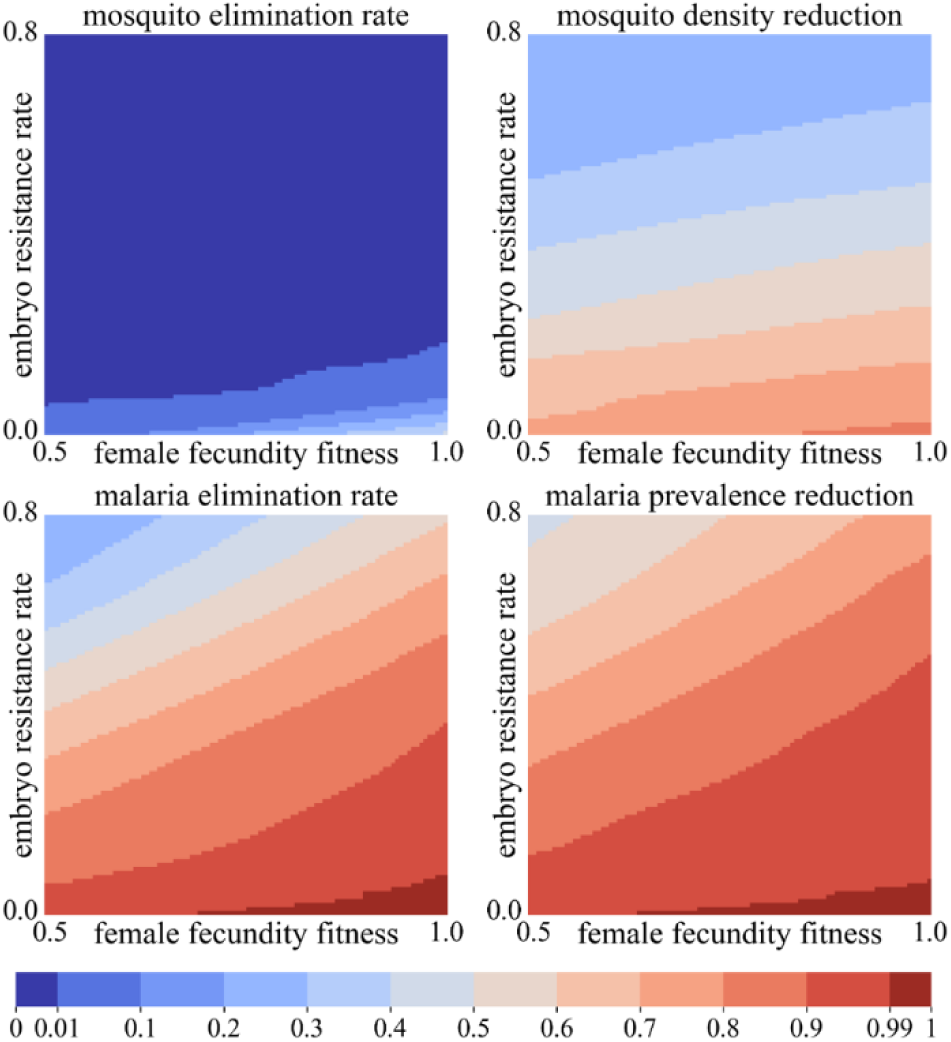
Effect of female heterozygote fitness and embryo resistance. Gene drive outcomes were predicted using the 3-layer deep learning model. Parameters are fixed at their default varies, with varying fecundity fitness of female drive heterozygotes and embryo resistance allele formation rate. We show the mosquito elimination rate, mosquito density reduction, malaria elimination rate, and malaria prevalence reduction compared to the pre-drive average.

**Figure S13.**
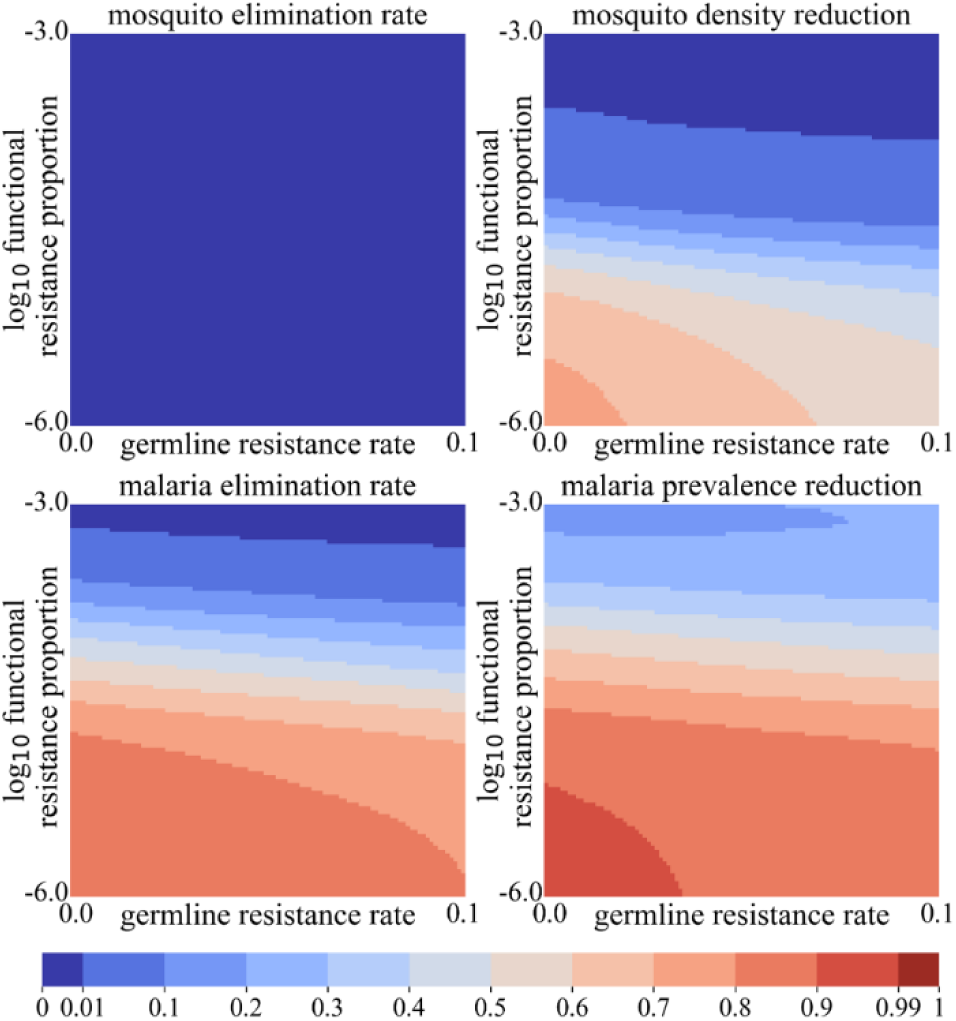
Effect of resistance allele formation. Gene drive outcomes were predicted using the 3-layer deep learning model. Parameters are fixed at their default varies, with varying germline resistance allele formation rate and relative proportion of functional resistance alleles. We show the mosquito elimination rate, mosquito density reduction, malaria elimination rate, and malaria prevalence reduction compared to the pre-drive average.

**Figure S14.**
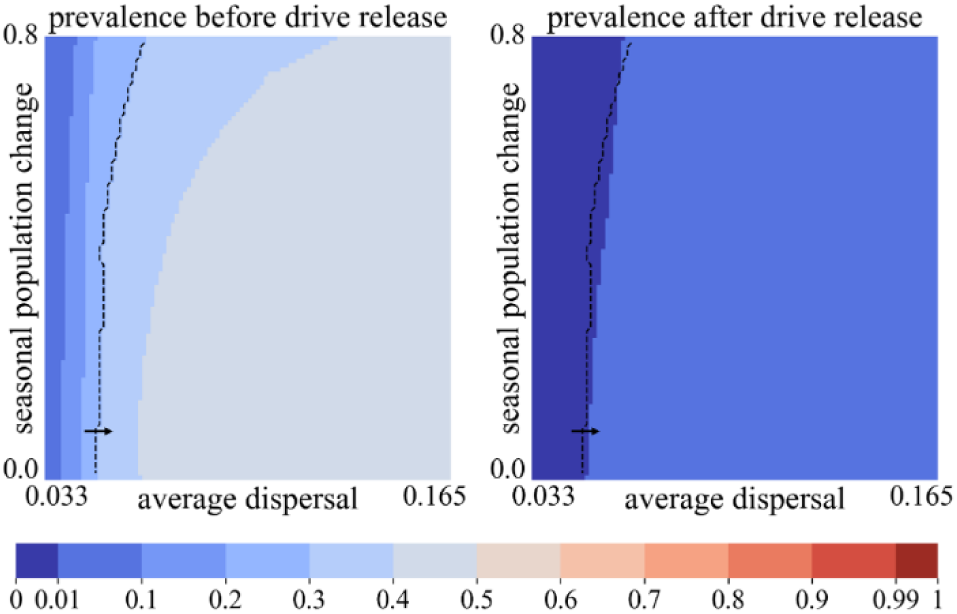
Effect of dispersal and seasonality. Gene drive outcomes were predicted using the 3-layer deep learning model. Parameters are fixed at their default varies, with varying average dispersal (km traveled by adults per week) and seasonal population change (increase and decrease, relative to average). We show the malaria elimination rate and malaria prevalence reduction compared to the pre-drive average. The deep learning model is trained using parameters with a malaria equilibrium prevalence in humans of between 30% to 50%. The arrow in the dashed lines shows where this parameter range is located on the heatmap (predictions outside this range may be less accurate).

## Notes

https://github.com/jchamper/Malaria-Drive-Modeling

### Competing Interest Statement

The authors have declared no competing interest.

